# Microbial hydrocarbon degradation potential of the Baltic Sea ecosystem

**DOI:** 10.1101/2025.06.02.657333

**Authors:** Joeselle M. Serrana, Benoît Dessirier, Francisco J. A. Nascimento, Elias Broman, Malte Posselt

**Affiliations:** Stockholm University Center for Circular and Sustainable Systems (SUCCeSS), Stockholm University, 106 91 Stockholm, Sweden; Department of Environmental Science (ACES), Stockholm University, 106 91 Stockholm, Sweden; Baltic Sea Centre, Stockholm University, Stockholm, Sweden; Department of Ecology, Environment, and Plant Sciences (DEEP), Stockholm University, 106 91 Stockholm, Sweden

**Keywords:** Baltic Sea, Benthic sediments, Environmental microbiome, Hydrocarbon degradation, Metagenomics, Pelagic water

## Abstract

**Background:** The Baltic Sea receives petroleum hydrocarbons from various point sources. The degradation of these contaminants in the environment is typically facilitated by a variety of microorganisms that possess a range of genes and metabolic functions related to the degradation of various hydrocarbon substrates. However, our understanding of natural attenuation and the microbial capacity to degrade these contaminants within the Baltic Sea ecosystem remains limited. In this study, we compiled metagenomes from the benthic and pelagic ecosystems across the Baltic Sea to identify microorganisms and characterize their genes and metabolic functions involved in the degradation of hydrocarbon compounds.

**Results:** Known hydrocarbon-degrading phyla, i.e., Pseudomonadota, Myxococcota A, Actinomycetota, and Desulfobacterota, were identified within the Baltic Sea metagenome-assembled genomes (MAGs). Notably, 80% of the MAGs exhibited multiple hydrocarbon degradation gene annotations (>10 reads per kilobase million). Aerobic degradation was the predominant pathway for hydrocarbon degradation across environmental samples. Hydrocarbon degradation gene abundances varied among samples and Baltic Sea subbasins, with long-chain alkanes and dibenzothiophene compounds being the preferred substrates. Species richness and diversity of both benthic and pelagic microorganisms positively correlated with hydrocarbon degradation gene diversity, with the pelagic ecosystem exhibiting significantly higher richness and diversity compared to the benthic ecosystem. Additionally, the composition of the hydrocarbon degradation genes across the Baltic Sea subbasins was influenced by oil spill history, with areas that experienced higher spill volumes showing lower microbial diversity, suggesting potential enrichment of specific hydrocarbon degraders. Among the environmental factors assessed, depth played a significant role in shaping the composition of genes involved in hydrocarbon degradation within the Baltic Sea.

**Conclusions:** Using metagenomics, we profiled the native microorganisms associated with hydrocarbon degradation in the Baltic Sea. This knowledge will aid in understanding the natural capacities of microbial communities, potentially linked to the natural attenuation of hydrocarbon pollutants in the area. Insights into microbial degradation potential can enhance predictions of petroleum pollutant persistence and accumulation, support mitigation strategies for marine pollution, and reveal the ecological resilience of native microbial communities in marine ecosystems.

**Graphical Abstract:** 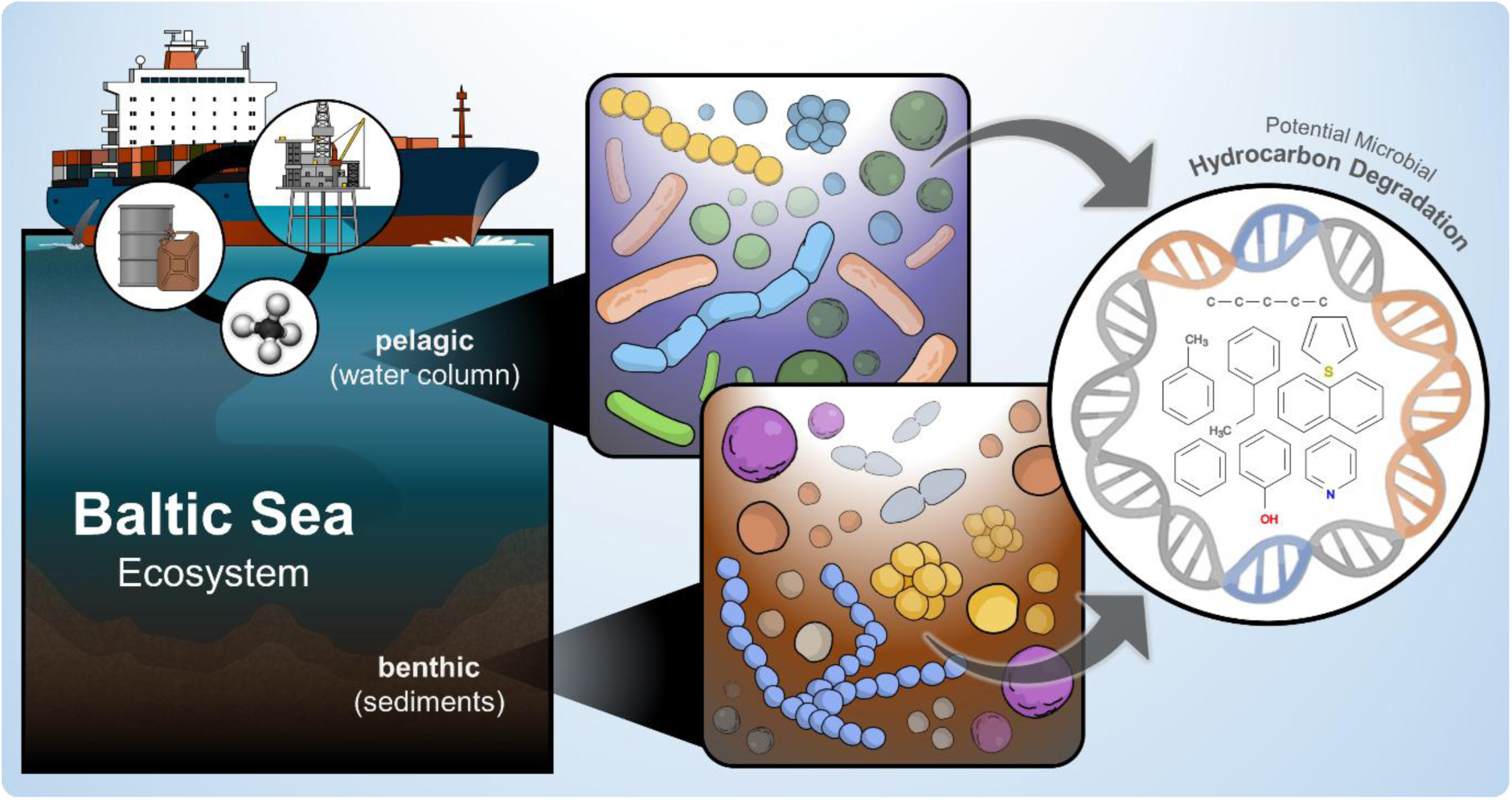

## Background

Petroleum hydrocarbons are one of the major organic pollutants in marine ecosystems [1,2]. These compounds disrupt ecological diversity and impair ecosystem functions, posing substantial threats to global health and sustainability [3–6]. While some hydrocarbons come from natural sources [7], more concerning issues arise from anthropogenic activities. This includes abrupt and large-scale inputs from oil spills, industrial discharges, atmospheric deposition, and marine oil and gas explorations [2,8,9], estimated to be 8.3 million tons introduced into marine ecosystems each year [10]. For instance, the Baltic Sea, one of the largest brackish water bodies, is also among the busiest waterways in the world [11,12]. Hazardous substances entering the Baltic catchment at elevated levels due to human activities remain among the most widespread and impactful pressures on the Baltic Sea [13,14]. The recent HELCOM holistic assessment for the 2016–2021 period (HOLAS 3) reported a continued overall decrease in both the frequency and volume of oil spills in the Baltic Sea, with accidental spills near port areas from cargo vessels or passenger ships rarely causing pollution [15].

Despite this, recent reports have documented negative impacts of oil spills and other petroleum products on the Baltic Sea’s marine and coastal ecosystems [12,16,17]. This is because the Baltic Sea basin remains vulnerable to the harmful effects of petroleum hydrocarbons and other hazardous substances [17,18]. These compounds can accumulate in sediments and marine organisms, leading to long-term ecological impacts, e.g., toxicity to marine life and disruption of local ecosystems [19,20]. Furthermore, potential future contamination continues to pose a risk due to various factors, e.g., increased maritime traffic and transportation, climate change, and the possibility of large-scale accidental spills [15]. For example, tankers traveling the Baltic Sea route have increased in both size and length, accompanied by growing shadow fleet activities [21–23]. This puts the area particularly at risk of oil spills and other maritime incidents. Therefore, while the situation has improved, ongoing monitoring and preventive measures remain critical to protect the Baltic Sea ecosystem.

The removal of petroleum hydrocarbons from the environment primarily depends on native microorganisms that degrade them by utilizing hydrocarbons as a source of carbon and energy [24]. The breakdown of these contaminants is generally facilitated by taxonomically diverse microorganisms that display a wide range of functions and substrates involved in hydrocarbon degradation [25–27]. Microbial degradation can occur under both aerobic and anaerobic conditions, with aerobic pathways employing oxygen to activate hydrocarbons and anaerobic pathways utilizing alternative electron acceptors, e.g., sulfate or nitrate, in oxygen-deprived environments [28]. Microbial degradation of hydrocarbons is a crucial ecosystem service vital for maintaining ecosystem health [26,29]. Assessing the natural microbial degradation of petroleum hydrocarbons is essential for understanding their environmental impacts on marine ecosystems and their inhabitants [28,30]. Previous studies have evaluated the ability of microorganisms to degrade hydrocarbons across various ecosystems [31–35]. For example, Miettinen et al. [35] studied the hydrocarbon degradation potential of bacterial, archaeal, and fungal microbiomes in the Baltic Sea, comparing long-term oil-exposed and less-exposed coastal sites. Using targeted gene identification (*alkB* and PAH ring hydroxylating deoxygenase) and taxonomic profiling via amplicon sequencing, they found similar microbial diversity and degradation potential at polluted and pristine sites in the Baltic Sea. However, our knowledge remains limited, as most research in marine ecosystems has focused on a few substrates or cultured microorganisms, resulting in a poorly characterized understanding of environmental microbiomes.

In this study, we profiled hydrocarbon-degrading microorganisms in both benthic and pelagic ecosystems of the Baltic Sea and characterized the associated genes and metabolic functions involved in the degradation of various hydrocarbon compounds. To the best of our knowledge, this is the first thorough profiling of the functional potentials of the Baltic Sea concerning hydrocarbon degradation. We utilized publicly available metagenomic datasets from environmental microbiomes sampled in the benthic [36,37] and pelagic [38,39] ecosystems across the Baltic Sea. Through this analysis, we identified potential microbial hydrocarbon degraders from both environments and screened for genes associated with the degradation of diverse hydrocarbon substrates. We also assessed the potential influence of environmental factors, i.e., water depth, salinity, and temperature gradients, on the degradation capacities of these environmental microbiomes. Additionally, we hypothesize that oil spills in the area have influenced the adaptive capacity and diversity of hydrocarbon-degrading microorganisms exposed to these contaminants. Hence, we investigated the potential link between the recorded annual average total oil spills in each subbasin of the Baltic Sea and their microbiomes’ ability to degrade hydrocarbons. Understanding the degradation potential of environmental microbiomes exposed to hydrocarbon contamination is essential for refining estimates of pollutant persistence and predicting the fate of petroleum hydrocarbons in marine and coastal environments. Our findings contribute to a deeper understanding of the functional resilience of Baltic Sea microbiomes and provide a scientific basis for developing improved management and mitigation strategies that address hydrocarbon pollution in aquatic ecosystems.

## Methods

### Compiling Baltic Sea metagenomes

A total of 203 Baltic Sea metagenomes from both benthic sediments (n = 108) [36,37] and water column samples (n = 95) [38,39] was compiled from public metagenomic data across four published studies (**Fig. 1a**). The benthic samples were collected from 2016 to 2019, while the pelagic samples were from a 2011 to 2015 collection. Sequence information and sample coordinates can be found in Supplementary Table S1. Environmental parameters, i.e., depth (m), salinity (PSU), and temperature (°C), compiled by Rodríguez-Gijón et al. [37], were utilized in this study. For further details on site description, sample collection, and sequencing, please refer to the respective publications of the public datasets. Following the HELCOM subdivisions of the Baltic Sea nomenclature, the metagenomes were assigned to Baltic Sea subbasins based on their location [40]. Two hundred samples were distributed across fifteen subbasins, while the remaining three samples were allocated to Skagerrak (referred to as SEA-000) (**Supplementary Fig. S1**).

**Fig. 1.**
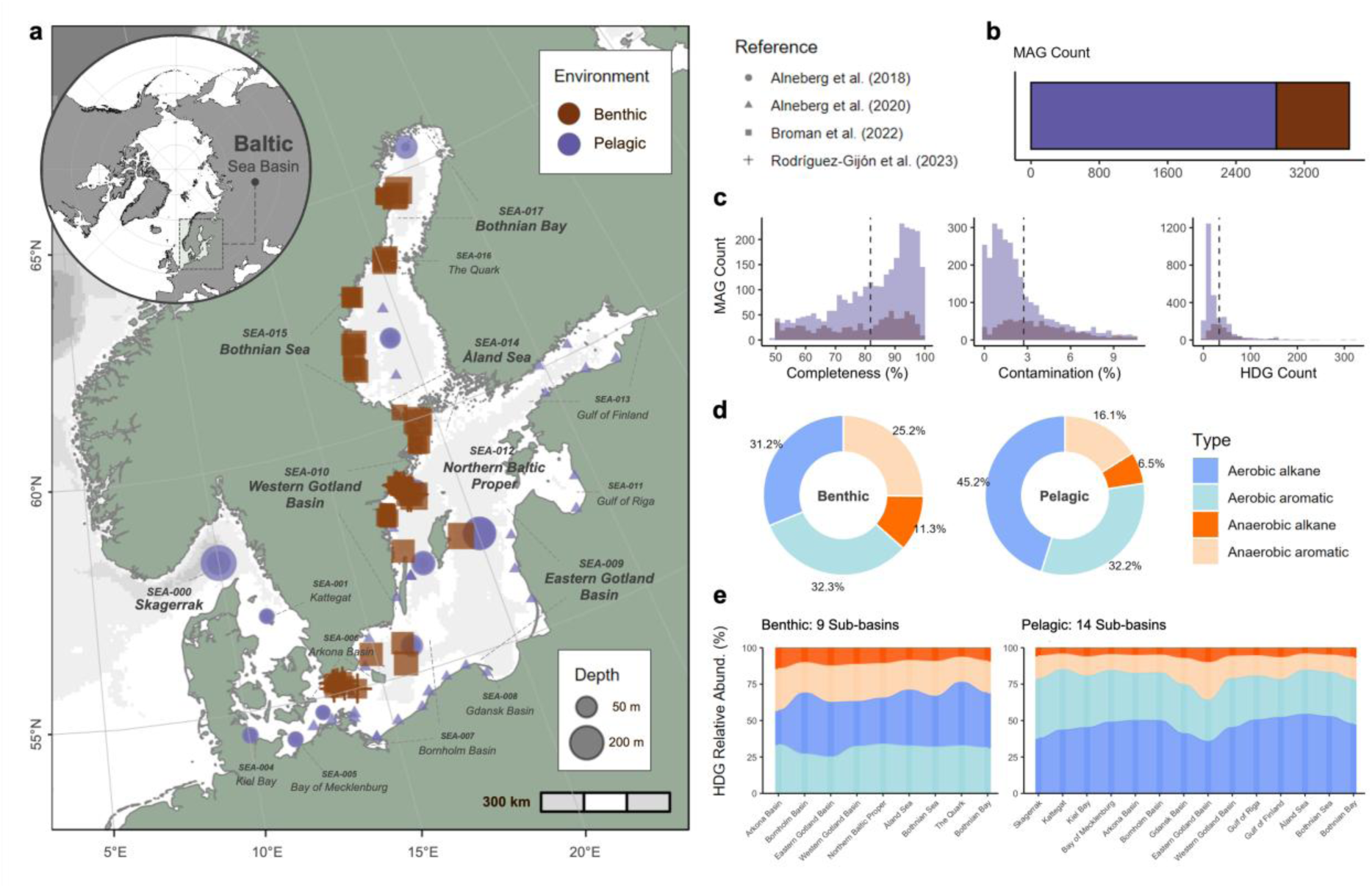
Baltic Sea metagenomics data and metagenome-assembled genome (MAG) information. (a) Location and sample environment of the metagenomics data compiled from Baltic Sea microbiome studies (i.e., Alneberg et al., 2018, 2020; Broman et al., 2022; Rodríguez-Gijón et al., 2023). (b) MAG counts and (c) histograms illustrating the distribution of MAG quality based on % completeness and % contamination and predicted hydrocarbon degradation gene (HDG) count. The relative abundance of HDGs by type based on (d) environmental samples and (e) across the Baltic subbasins.

### Metagenomic assembly, binning, and phylogenomic analysis

The raw paired-end sequences of the 203 metagenomic datasets were adapter-trimmed and quality-filtered using fastp v0.23.4 [41], followed by the removal of human and PhiX spike-in reads with Kraken2 v2.1.2 [42]. The quality-filtered reads were then error-corrected using bbcms version 38.61b from BBTools [43], with parameters set to a minimum count of 2 and a high-count fraction of 0.6. Individual assemblies of the 203 quality-filtered metagenomic datasets were conducted with MEGAHIT v1.2.9 [44]. Genes were then predicted on contigs using Prodigal v2.6.3 [45], employing the -p meta option. The read counts for each predicted gene were estimated by mapping the quality-filtered paired-end reads to the contigs using Minimap2 v2.24-r1175 [46] and summarizing the read abundances into count tables with featureCounts v2.0.3. [47] Read processing counts at each step are listed in **Supplementary Table S2**.

Binning was performed using the single_easy_bin module of SemiBin v2.1 [48]. Binning parameters were set to default with the environment flag set to global. The bins were then evaluated for contamination and completeness using CheckM v1.1.3 [49] with lineage_wf and default parameters. A total of 10,017 bins were recovered from the individual assemblies. From this, 13.6% (1,318) were classified as high-quality (completeness of >90% and contamination of <5%), while 37.2% (3,726) were medium-to-high quality (completeness of >50% and contamination <10%) (**Supplementary Table S3** and **Fig. 1a-b**). The 193 samples with medium-to-high quality bins were retained in the downstream analysis, as their inclusion broadened the dataset, enabling a more extensive exploration of microbial capabilities and referred to as metagenome-assembled genomes (MAGs). The average guanine and cytosine (GC) base content was 52%, with an average contig (N50) length of 19,833 and a predicted gene count of 2,230, respectively. The average genome size was 2.20 million base pairs (Mbp), with the largest measuring 8.89 Mbp (ERR2206771.bin_3) (**Supplementary Fig. S2**).

Taxonomic assignment was performed on the MAGs using GTDB-Tk v2.4.0 with R220 [50] through the classify_wf function. The MAG quality parameters and taxonomic information are presented in **Supplementary Table S3**. The protein sequence alignments produced from GTDB-Tk were subsequently trimmed using trimAl v1.4.rev15 [51] with the heuristic “-automated1” method and were used as inputs to construct phylogenomic trees with IQTree2 multicore v2.2.0.3 COVID-edition [52]. The resulting trees were visualized and annotated using the Interactive Tree Of Life (iTOL) v4.4.2 [53].

### Screening of hydrocarbon degradation genes

Hydrocarbon degradation genes were screened using the Calgary approach to ANnoTating HYDrocarbon degradation genes (CANT-HYD) [54]. Genes involved in hydrocarbon degradation were identified using Hidden Markov Model (HMM) profiles from the CANT-HYD database, with a noise cut-off E-value of 0.01 for the binned metagenomes. This approach annotates 37 key marker genes associated with anaerobic and aerobic degradation pathways for both aliphatic and aromatic hydrocarbons, based on experimental and in silico-inferred functional enzymes (**Supplementary Table S4**). Gene abundances were expressed as reads per kilobase million (RPKM) to normalize gene coverage values, correcting for differences in sample sequencing depth and gene length [55]. The RPKM counts for each gene per sample are provided in **Supplementary Table S5**.

### Statistical analysis

All visualizations and statistical analyses were conducted using R v4.4.3 [56]. A Pearson correlation analysis was used to assess the relationship between genome size and the number of annotated hydrocarbon degradation genes (HDGs). The microeco v1.13.0 package [57] was utilized to calculate both alpha (i.e., Observed, Chao1 richness, Shannon diversity, and Pielou evenness) and beta diversities (i.e., Jaccard dissimilarity index) of the microbial community and HDG datasets. A t-test or a one-way analysis of variance (ANOVA) with Duncan’s multiple comparison tests was employed to examine significant differences in alpha diversity across categories. Principal coordinate analysis (PCoA) was performed to visualize the Jaccard dissimilarity matrices, and a non-parametric multivariate analysis of variance (PERMANOVA) was utilized to infer significant differences in composition for both the taxonomic and HDG datasets. A Pearson correlation analysis was conducted to investigate the relationships between species and HDG diversity indices, and to assess the potential connection between microbial diversity and HDG composition. A random forest analysis coupled with a differential abundance test [58] was performed on the HDG dataset to identify differentially abundant HDGs between environments. The annual average total oil spill volumes during the assessment period of 2016-2021 (m³), as reported by HELCOM [15] (**Supplementary Table S6**), were used to correlate oil spills in each Baltic Sea subbasin with their hydrocarbon degradation gene profile. A Spearman’s correlation analysis was conducted to assess the relationships between environmental factors and HDGs using the microViz v0.12.1 package [59]. Skagerrak samples were excluded from this analysis since the area is not covered in the oil spill reports recorded by the commission.

## Results

### Microbial hydrocarbon degraders from the benthic and pelagic ecosystems

We recovered 3,726 medium-to-high quality MAGs with an average genome size of 2.20 Mbp, a completeness of 81.8%, and 2.7% contamination (**Fig. 1c**). Of these, 854 MAGs were identified in benthic samples, while 2,872 MAGs were found in pelagic samples. The average genome size of benthic MAGs was 2.73 Mbp, whereas the pelagic MAGs had 2.04 Mbp (**Supplementary Table S7**). The difference in average genome sizes between the environments may be linked to the distinct evolutionary pressures and ecological roles these microorganisms fulfill within their respective habitats.

Using a validated HMM profiling approach, genes encoding hydrocarbon-degrading enzymes were identified in MAGs. Most of these, except for a bacterial MAG (ERR2206774.bin_159) and an archaeal MAG (ERR2206790.bin_40), had at least one hydrocarbon degradation gene (HDG). All 37 key HDGs were identified from the compiled Baltic Sea metagenomic samples. Multiple copies of HDGs can be found within a single MAG, with an average of 34 HDG reads per kilobase million (RPKM) (Fig. 1c). A total of 213 MAGs had >100 HDGs annotated. For example, one bacterial MAG (ERR5010709.bin_8) had 320 RPKM, which had the highest HDG annotation. Benthic MAGs had higher HDG annotations, with an average of 49 RPKM, compared to 29 in pelagic MAGs. We found a statistically significant but moderately positive linear relationship between genome size and HDG count for all MAGs (r = 0.43, t = 29.39, df = 3,724, p < 0.001) (**Supplementary Fig. S3**). By environment, the genome size of benthic MAGs showed a stronger relationship with HDG count (r = 0.59, t = 21.424, df = 852, p-value < 0.001) compared to the pelagic samples (r = 0.35, t = 19.852, df = 2870, p< 0.001). Based on four metabolic categories, the hydrocarbon degradation potential of benthic and pelagic MAGs differed across the assessed pathways. HDGs under the aerobic alkane metabolism category were notably higher in the pelagic MAGs (45.2%) compared to benthic MAGs (31.2%) (**Fig. 1d**). Conversely, HDGs involved in anaerobic alkane metabolism were more pronounced in the benthic MAGs (11.3%) than in the pelagic MAGs (6.5%). For the aromatic degradation pathways, HDGs involved in anaerobic metabolism were higher in benthic MAGs (25.2%) compared to pelagic (16.1%), while HDGs for aerobic metabolism exhibited similar levels, with 32.3% in benthic and 32.2% in pelagic MAGs.

Moreover, the relative abundance of the four types for each subbasin is shown in **Fig. 1e**. Based on the frequency of HDG annotation, each subbasin exhibited varying levels of abundance (**Supplementary Fig. S4**), with the Western Gotland Basin samples (SEA-010) having the highest number of identified HDGs. In support of this, the observed and HDG Chao1 richness and Shannon diversity of each subbasin in both environments were significantly different (p < 0.01) (**Supplementary Fig. S5**). In contrast, Pielou’s evenness index was not significantly different among subbasins for both environments (benthic, p = 0.78; pelagic, p = 0.08). According to the HMM annotations, 25 hydrocarbon degradation marker gene annotations were identified across all Baltic Sea subbasins (**Supplementary Fig. S6**).

Of the MAGs with hydrocarbon degradation potential, 79 were archaeal genomes (**Supplementary Fig. S7**), while 3,645 were classified as bacteria according to the GTDB-tk classification (**Fig. 2a**). All MAGs had classifications at least up to the order level. Three MAGs had unclassified families, and 146 MAGs lacked classified genera. In total, 2,526 MAGs (67.8%) had species-level identifications (**Fig. 2b**). The MAGs represented a diverse array of prokaryotes, encompassing 35 phyla (5 archaeal and 30 bacterial), 243 families (9 archaeal and 234 bacterial), 467 genera (15 archaeal and 452 bacterial), and 657 species (22 archaeal and 634 bacterial, including unclassified species). For the archaeal annotations, the MAGs showed that 67.9% (53) were Thermoproteota, followed by Thermoplasmatota (26.9%), Nanoarchaeota (2.6%), Micrarchaeota (1.3%), and Halobacteriota (1.3%). In the bacterial phyla, MAGs assigned to Pseudomonadota (33.6%) were the most prevalent, followed by Actinomycetota (22.3%), Bacteroidota (14.9%), Planctomycetota (5%), and Desulfobacterota (4.9%). The remaining bacterial phyla accounted for less than 5% of the MAGs. The relative abundance of the top 10 phyla across each environment is presented in **Fig. 3a**. Pseudomonadota predominated most samples, except for the benthic Eastern Gotland Basin, where the MAGs were mostly assigned to Gemmatimonadota. The relative abundance of Desulfobacterota in benthic samples decreased from the Arkona and Bornholm Basins to the more freshwater subbasins, e.g., Åland Sea, Bothnian Sea, and Bothnian Bay. Conversely, Actinomycetota exhibited an increasing trend from the more marine subbasins, e.g., Skagerrak and Kattegat, to freshwater sites.

**Fig. 2.**
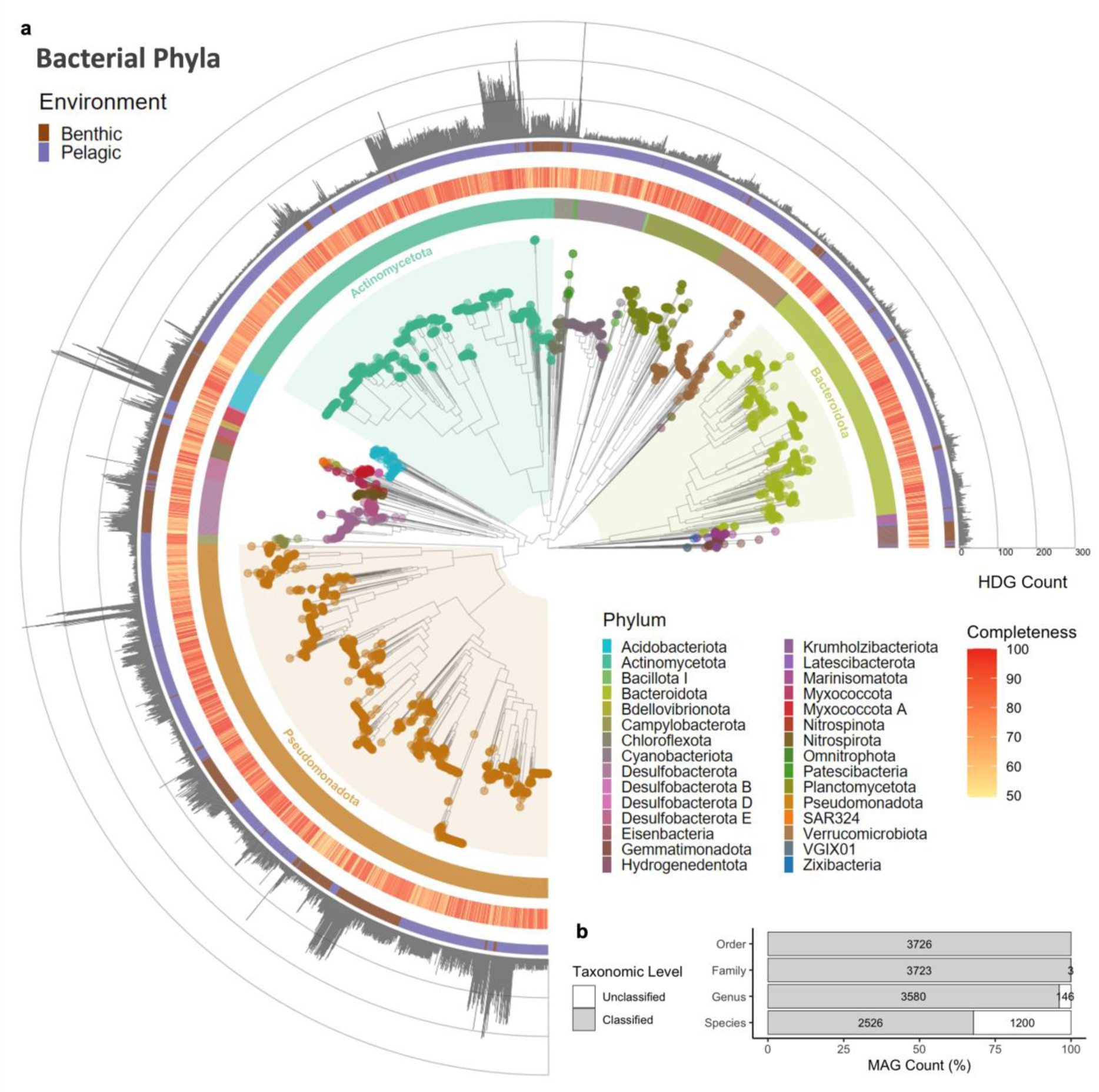
Taxonomic assignment of MAGs. (a) Maximum-likelihood phylogenomic tree of bacterial MAGs recovered from Baltic Sea metagenomic samples. (b) Percentage of MAGs assigned at various taxonomic ranks based on GTDB-tk classification.

**Fig. 3.**
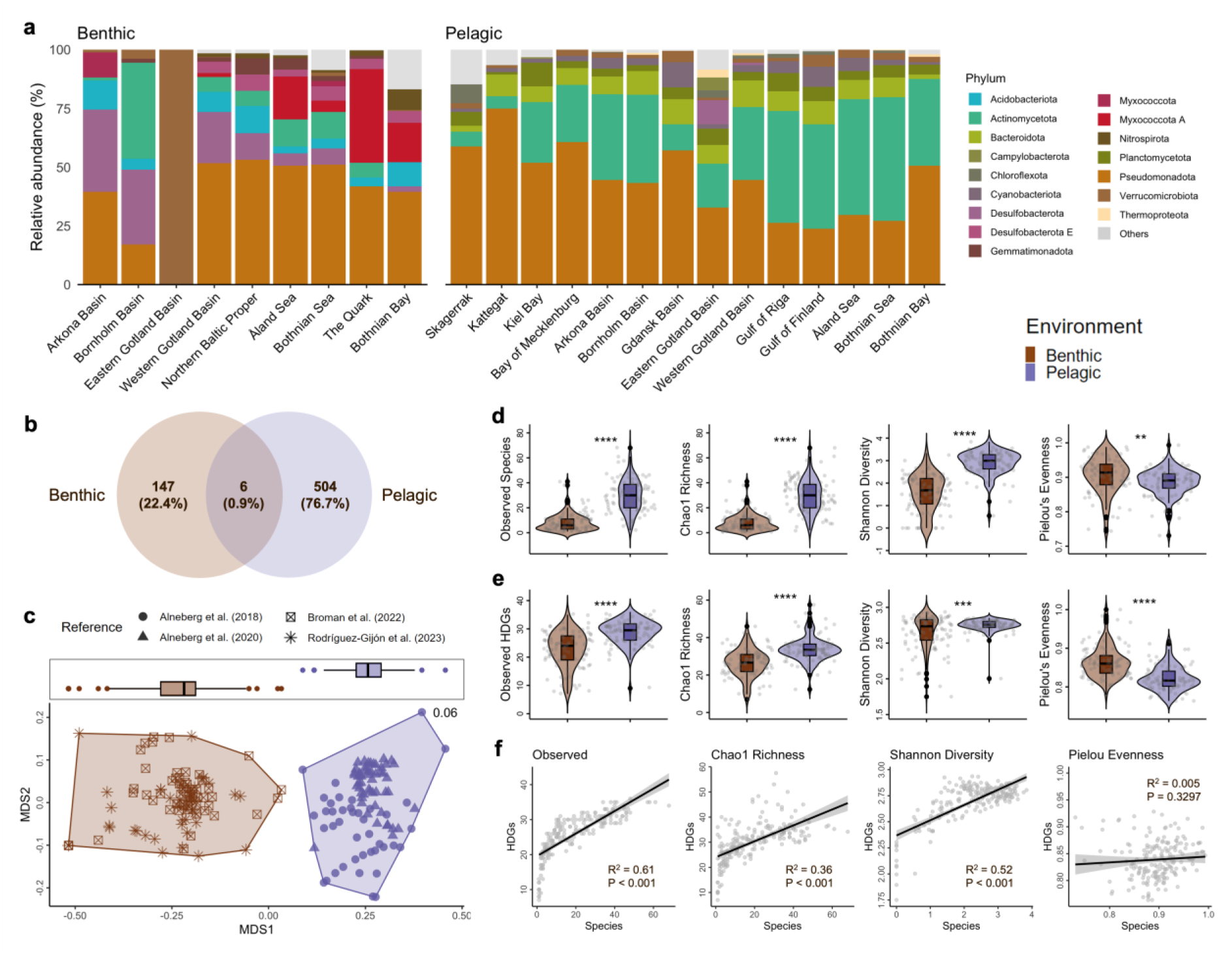
Taxonomic and hydrocarbon degradation gene (HDG) composition and diversity. (a) Relative abundance of the top microbial phyla in each sub-basin and environment. (b) Shared and unique species per environment. (c) Non-metric multidimensional scale (NMDS) plot of species composition based on Jaccard dissimilarity. (d) Alpha diversity estimates of the (e) species and (f) HDGs. “****” indicates differences between groups tested by t-test at p < 0.001, “***” at p < 0.01.

At the species level, only six species (0.9%) were found to be present in both environments (**Fig. 3b**). Including MAGs with unclassified species annotations, the benthic samples comprised 147 species, whereas the pelagic samples included 504 species. The most abundant species for each environment are depicted in **Supplementary Fig. S8**. The species with the most annotations was *UBA1847 sp022866645* (Pseudomonadota, Woeseiaceae) with a total of 61 MAGs (1.64%), found exclusively in benthic samples, followed by *D2472 sp002358345* (Pseudomonadota, D2472) with 59 MAGs (1.58%), *MAG-120802 sp030730425* (Actinomycetota, Nanopelagicaceae) with 53 MAGs (1.42%), and *UBA9145 sp001438145* (Pseudomonadota, Pseudohongiellaceae) with 52 MAGs (1.40%), which were only found in pelagic samples. The counts and total relative abundance of all species are listed in **Supplementary Table S9**, and a visualization of the shared and unique species per sub-basin is presented in **Supplementary Fig. S9**. Three MAGs identified as *JAJDAM01 sp.* (Myxococcota A, SMWR01) (i.e., ERR5010709.bin_8, ERR5010720.bin_14, and ERR5010722.bin_46), and two *CALGIA01 sp937916005* (Pseudomonadota, MPNO01) had multiple HDG annotations > 300 RPKM (**Supplementary Table S3**).

The species composition of the two environments was distinctly separated from one another based on Jaccard dissimilarity (**Fig. 3c**). PERMANOVA revealed significant effects of both the environment and subbasin on species composition. The environment explained 11.87% of the variance (R² = 0.12, F = 29.36, p = 0.001), while the subbasin accounted for 16.94% (R² = 0.17, F = 2.79, p = 0.001). The analysis of similarities (ANOSIM) confirmed strong differentiation between the pelagic and benthic samples (R = 0.69, p = 0.001). The homogeneity of multivariate dispersion (betadisper) test also indicated no significant differences in dispersion among the groups (F = 2.35, p = 0.137), supporting the assumption of equal variances across the groups. Pairwise t-tests further revealed significant differences in group distances between benthic and pelagic communities (p.adj < 0.001), suggesting a clear separation between these groups (**Supplementary Table S9**). Additionally, the alpha diversity comparison of the two environments further supports the differentiation between the benthic and pelagic samples (**Fig. 3d**). The observed species (30 ± 1 vs. 9 ± 7), Chao1 richness (30 ± 1 vs. 9 ± 7), and Shannon diversity (2.9 ± 0.5 vs. 1.6 ± 0.8) in the pelagic environment were significantly higher than in the benthic environment (p.adj < 0.001). Moreover, Pielou’s evenness metric (0.99994 ± 0.0003 vs. 0.99997 ± 0.0002) demonstrated a significant difference in the evenness of the microbial taxa between environments.

### Genes and metabolic pathways associated with the degradation of various hydrocarbon substrates

The top ten most abundant HDGs found in the benthic samples included the dibenzothiophene desulfurization enzyme C [*dszC (soxC)*], the flavin-binding monooxygenase (*alma Group I* and *III*), the putative alkane C2 methylene hydroxylase (*ahyA*), ethylbenzene dehydrogenase (*ebdA*), p-cymene dehydrogenase (*cmdA*), naphthalene dioxygenases (*non ndoB type* and *ndoB*), the monoaromatic dioxygenase (*bphA/tcbA/ipbA/bnzA*), and the long-chain alkane monooxygenase (*ladB*) (**Fig. 4a**). Similarly, except for the *cmdA* gene, these HDGs were also abundant in the pelagic samples with the addition of the cytochrome P450 alkane hydroxylase (*cyp153*). The occurrence of each HDG per phylum is presented in Supplementary Fig. S10. Pseudomonadota had the most prevalent and abundant HDGs, followed by Myxococcota A, Chloroflexota, Actinomycetota, and Desulfobacterota.

**Fig. 4.**
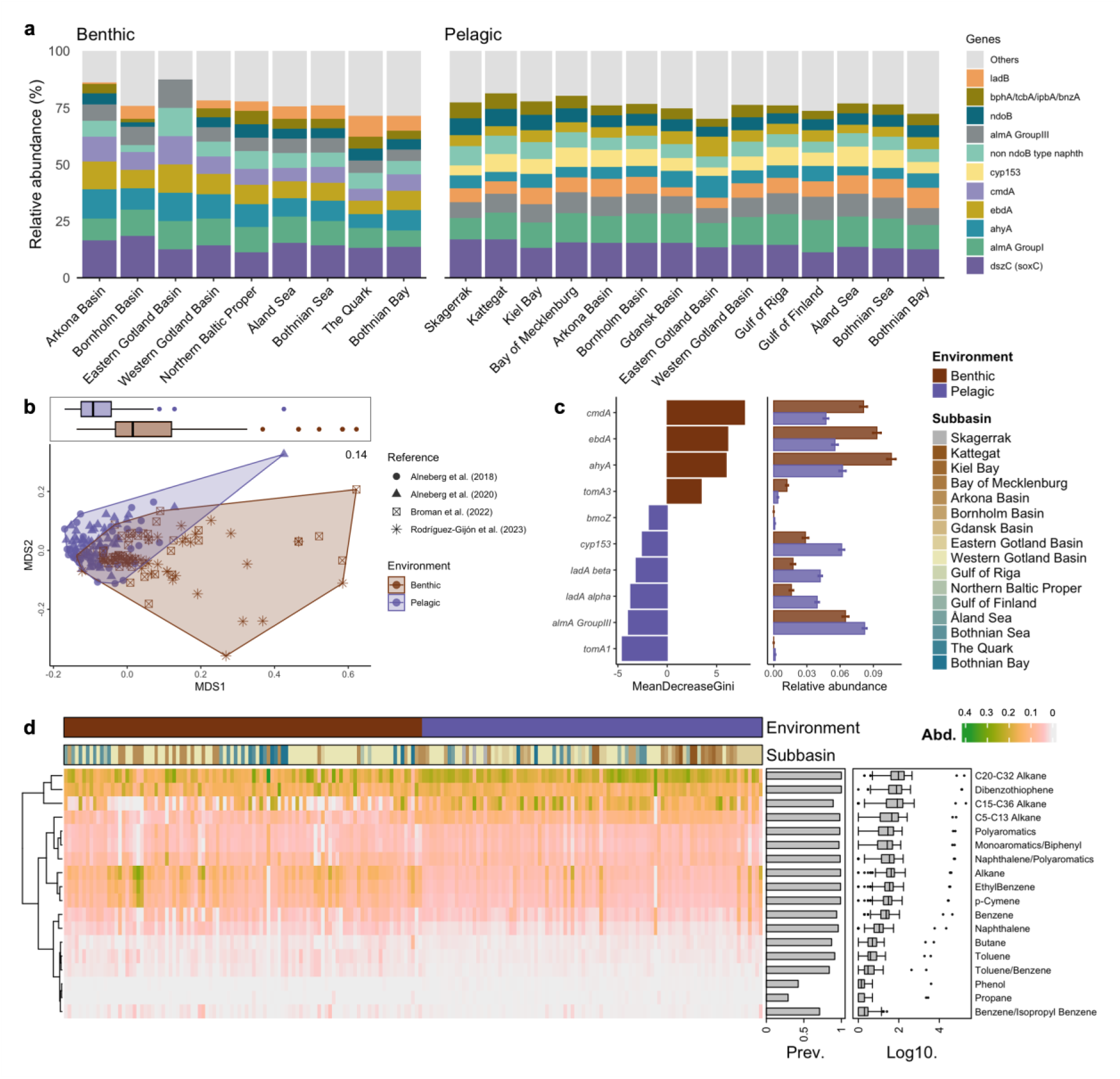
Hydrocarbon degradation genes (HDGs) and their substrates. (a) Relative abundance of the top HDGs in each sub-basin and environment. (b) Non-metric multidimensional scaling (NMDS) plot of HDG composition based on Jaccard dissimilarity. (c) Random forest classification of the relative abundance of HDGs across environments: (left) HDGs with Mean Decrease Gini > 2, presented in descending order of importance; see Supplementary Fig. S11 for the plot of all significantly differential genes; (right) Relative abundance of HDGs enriched in each environment. The bars represent the mean relative abundance, and error bars indicate the standard error. (d) Heatmap of the relative abundance of hydrocarbon substrates associated with the degradation genes. Hierarchical clustering analysis was performed using Ward’s method and Euclidean distance metric.

The alpha diversity estimates of the HDGs displayed similar trends to species alpha diversity, with pelagic samples showing higher diversity and richness values compared to benthic samples, and a significantly greater evenness in benthic samples (**Fig. 3e**). Interestingly, the correlation between their alpha diversity values (**Fig. 3f**) revealed a strong positive relationship between observed species and HDGs (R² = 0.61, p < 0.001), indicating that as HDGs increased, the number of observed species also increased significantly. Similarly, the Shannon diversity index showed a moderate positive correlation with HDGs (R^2^ = 0.515, p < 0.001), suggesting a link between increasing HDGs and species richness and evenness. In contrast, the species Chao1 richness exhibited a weaker correlation with HDGs (R^2^ = 0.358, p < 0.001). Notably, Pielou’s evenness demonstrated a negligible correlation with HDGs (R^2^ = 0.005, p < 0.001), indicating that the species had minimal influence on the observed HDG evenness.

For the beta diversity of HDG composition, Jaccard dissimilarity revealed differences between the benthic and pelagic samples (**Fig. 4b**). PERMANOVA demonstrated significant effects from both environmental factors and subbasin grouping on the HDG composition. The environment accounted for 15.13% of the variance (R² = 0.15, F = 38.17, p = 0.001), while subbasin explained 15.12% (R² = 0.1512, F = 2.54, p = 0.001). These findings were further supported by ANOSIM, which indicated moderate differentiation among the environmental samples (R = 0.24, p = 0.001). A betadisper test also showed significant differences in dispersion among groups (F = 6.73, p = 0.01), with pairwise comparisons indicating differences between benthic and pelagic samples (p = 0.01). Finally, the group distance comparison revealed highly significant differences in HDG composition between benthic and pelagic samples (p.adj < 0.001), emphasizing their distinct separation. These results highlight the crucial roles of the environment in shaping HDG compositional patterns (**Supplementary Table S9**). Random forest analysis identified 34 hydrocarbon degradation marker gene annotations significantly associated with specific environments. (**Fig. 4c**, **Supplementary Fig. S11** and **Table S10**). In the benthic samples, *cmdA*, *ebdA*, and *ahyA* exhibited the highest importance scores (Mean Decrease Gini values) of 7.85, 5.98, and 5.89, respectively. In the pelagic samples, t*omA1*, *almA Group III*, and *ladA alpha* showed high influence with scores of 4.61, 4.37, and 3.51. These differences, supported by strong statistical significance (P.adj < 0.001), emphasize niche specialization and its impact on shaping microbial dynamics and hydrocarbon degradation functions in each environment.

### Substrate preferences of potential hydrocarbon degraders

**Fig. 4d** and **Supplementary Table S8** show the distribution of substrates associated with the HDGs identified in benthic and pelagic samples. Among the substrates, genes linked to C20-C32 alkanes display the highest abundance, contributing 18.67% of the total RPKM HDG abundance, with a significant presence in both benthic (6,548) and pelagic (16,927) zones. This is followed by genes related to C15-C36 alkanes (14.86%) and dibenzothiophene degradation (14.24%), which are also predominantly found in pelagic environments. Genes involved in the degradation of short-chain alkanes (C5-C13) make up 8.80% of the total, while those associated with generic alkane degradation account for 7.09%. Other notable contributions come from genes related to naphthalene/polyaromatics (6.38%), ethylbenzene (6.04%), and polyaromatics (5.59%). Minor contributions are observed for compounds like benzene (3.63%), naphthalene (1.88%), and toluene (0.78%), with trace levels of genes associated with propane (0.07%) and phenol degradation (0.11%). We emphasize the prevalence of genes associated with the degradation of long-chain alkanes and certain aromatic compounds in Baltic Sea environments, with pelagic zones typically exhibiting higher gene abundances than benthic zones.

Using a random forest analysis to assess the differential abundance of genes between the environments, genes involved in the degradation of the substrates p-Cymene, alkanes, and ethylbenzene exhibited the highest importance values of 12.21, 8.63, and 8.60, respectively, which were differentially abundant in benthic samples (p < 0.0001). Other compounds, i.e., toluene/benzene, naphthalene, and benzene, also contributed to the classification, albeit to a lesser extent (Mean Decrease Gini values > 3). In the pelagic environment, the genes associated with the degradation of long-chain (i.e., C15-C36 and C20-C32) and short-chain alkanes (i.e., C5-C13) were the most differentially abundant, with values of 5.04, 4.33, and 3.52 (p < 0.001). Additionally, phenol and polyaromatics demonstrated notable importance, while monoaromatics/biphenyl and propane also contributed to the random forest classification model (**Supplementary Fig. S12** and **Table S10**). We also observed interesting patterns based on the relative abundance of the substrates degraded by the HDGs across the Baltic Sea. Although the HDGs associated with the degradation of C20-C32 alkanes were the most abundant substrates in each sub-basin, the Gulf of Finland stands out for having the highest relative abundance (24.2% of the HDGs identified in the sub-basin), suggesting elevated activity for degrading these substrates in this area, followed by the Gulf of Riga (23%) and the Bay of Mecklenburg (22%). The subbasins Skagerrak (8%) and Kattegat (8%) exhibited relatively higher relative abundances of polyaromatics compared to other areas (**Supplementary Table S10**).

### Correlations among hydrocarbon degradation potential, environmental variables, and oil spill volume

The annual average of total oil spills across the subbasins during the 2016-2021 assessment period varied significantly (**Supplementary Table S1** and **Fig. 5a**). The Gulf of Finland recorded the highest average spill volume at 3.10 m³, followed by the Northern Baltic Proper at 1.71 m³ and the Kattegat at 1.31 m³. Moderate spill levels were observed in subbasins such as the Arkona Basin (1.23 m³), Bornholm Basin (1.08 m³), and Western Gotland Basin (0.38 m³). The Gulf of Riga and the Quark reported negligible or no recorded oil spills [15]. In addition to depth, salinity, temperature, latitude, and longitude (**Supplementary Table S1**), we evaluated the relationship between the HDG composition and annual average total volume of oil spills.

**Fig. 5.**
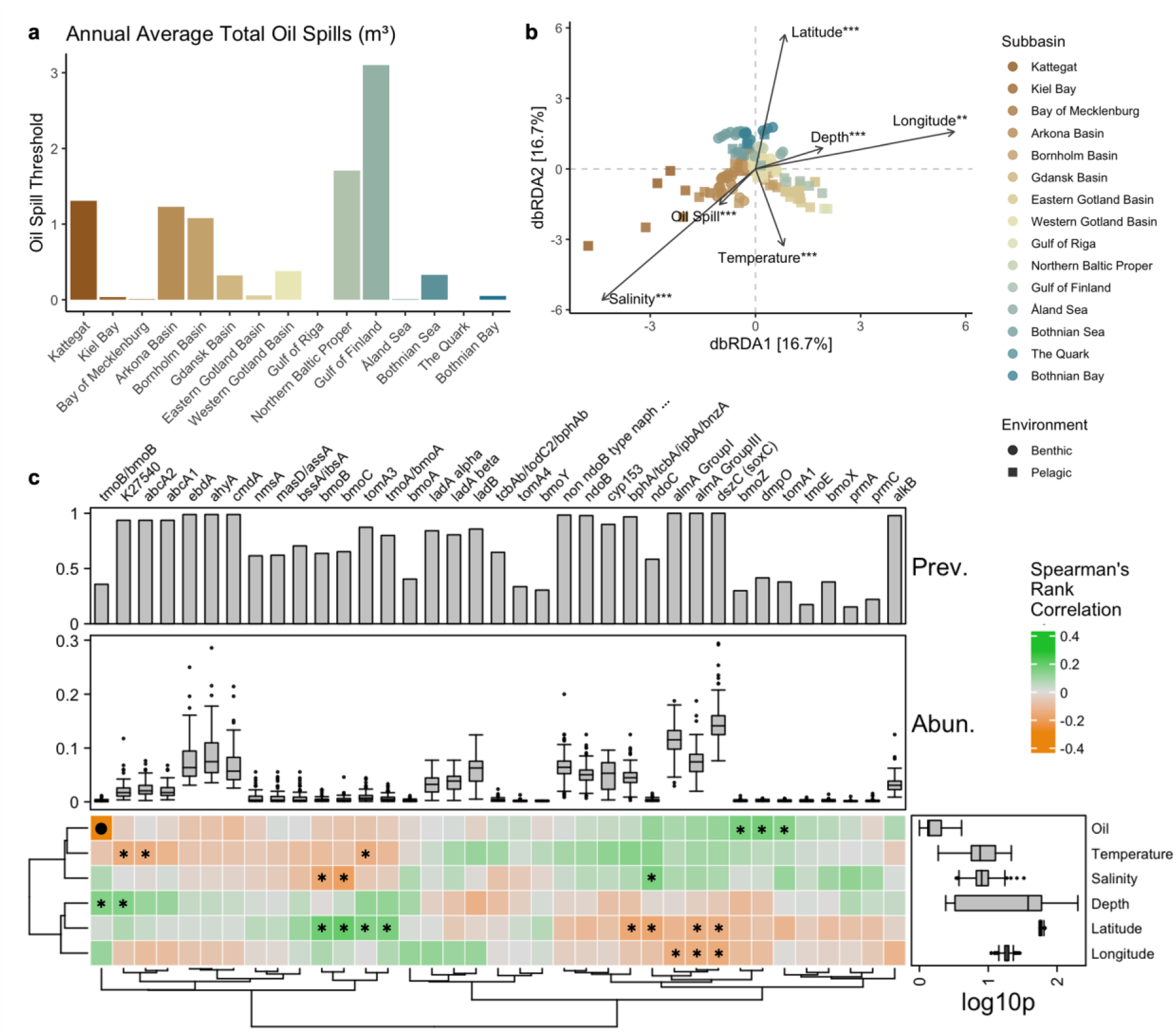
Environmental and oil spill association of the hydrocarbon degradation gene (HDG) profile. (a) The annual average total volume of oil spills (m³) for each subbasin (excluding Skagerrak) from 2016 to 2021, as reported by HELCOM (2023b). (b) Distance-based redundancy analysis (dbRDA) ordination plot of the HDG composition based on Jaccard distance and the environmental variables. “***” indicates a significant p < 0.001, “**” p < 0.01, and “*” p < 0.05. (c) Spearman’s rank correlation illustrating the relationship between the HDGs and environmental variables. Positive correlations are indicated in green, while negative correlations are shown in orange. The asterisk indicates p < 0.05, and the filled circle represents FDR-corrected p < 0.05. “Oil” refers to total oil spills (m³). Hierarchical clustering analysis was conducted using Ward’s method and the Euclidean distance matrix.

Distance-based redundancy analysis (dbRDA) revealed significant relationships between environmental variables and multivariate response patterns. Depth was the most influential factor, accounting for 80.27% of the variance (R² = 0.80, p < 0.001). Other significant variables included the oil spill threshold (R² = 0.63, p < 0.001), salinity (R² = 0.28, p < 0.001), and latitude (R² = 0.18, p < 0.001). In contrast, longitude and temperature, although significant (p < 0.005), explained smaller proportions of the variance (R² = 0.05 and R² = 0.15) (**Fig. 5b** and **Supplementary Table S11**).

Furthermore, Spearman’s rank correlation analysis (**Fig. 5b** and **Supplementary Table S12**) offered statistically significant insights into the relationships between each environmental variable and the specific HDGs. For example, the correlation between depth and the genes *K27540* (estimate: 0.147, p-value < 0.05) and *tmoB/bmoB* (estimate: 0.147, p-value < 0.05) indicates weak positive associations, suggesting that the expression levels of these genes may increase slightly with depth. Similarly, latitude showed mixed relationships across various genes. Positive correlations, e.g., with *bmoC* (estimate: 0.199, p-value < 0.05), suggest that these gene may increase with latitude, whereas negative correlations, e.g., with *ndoC* (estimate: −0.174, p-value < 0.01) and *almA Group III* (estimate: −0.147, p-value < 0.05), imply reduced expression at higher latitudes. The *bmoZ* (estimate: 0.172, p-value < 0.05), *dmpO* (estimate: 0.017, p-value < 0.05), and *tomA1* (estimate: 0.014, p-value < 0.05) genes were positively correlated with the annual average total oil spills. This suggests that the expression of this hydrocarbon degradation gene increases with increasing oil volume. On the other hand, the relationship between oil spill volume and *tmoB/bmoB* is particularly notable, demonstrating a strong negative correlation (estimate: −0.291, p-value < 0.001, FDR: 0.011). While these correlations revealed several potentially meaningful interactions between environmental variables and HDGs, including the strong and significant link between oil spill and *tmoB/bmoB*, most of the other associations were statistically not significant after controlling for multiple tests (i.e., adjusted FDR), which warrants cautious interpretation of the weaker trends.

## Discussion

### Most Baltic Sea MAGs have the potential to degrade hydrocarbons

Pseudomonadota was the most prevalent phylum and had multiple HDG annotations, followed by Myxococcota A, Actinomycetota, and Desulfobacterota. These phyla were identified as broad-spectrum hydrocarbon-degrading bacteria from the global ocean metagenome.^35^ In particular, Pseudomonadota comprises several species renowned for their ability to degrade hydrocarbons, including *Pseudomonas spp.* and *Acinetobacter spp.*^60,61^ These bacterial species can degrade a wide range of hydrocarbons, e.g., alkanes, aromatic compounds, and polycyclic aromatic hydrocarbons, often using them as their sole carbon source [60]. The degradation process involves specialized enzymes, e.g., oxygenases, which activate and transform most harmful hydrocarbons (e.g., cyclic and aromatic hydrocarbons) into less toxic compounds (e.g., linear alkanes) [60,62]. This makes Pseudomonadota crucial for bioremediation efforts in environments contaminated by petroleum and other hydrocarbon pollutants [63,64]. In addition, Myxococcota A, Actinomycetota, and Desulfobacterota were previously identified as hydrocarbon degraders in petroleum-rich marine or brackish sediments [65–68].

We identified all 37 key marker genes linked to anaerobic and aerobic degradation pathways for both aliphatic and aromatic hydrocarbons [54]. Eighty percent of the medium-to-high quality MAGs recovered had at least 10 HDG annotations and exhibited multiple copies within a single MAG, with an average HDG count of 34 RPKM. Additionally, 5.7% of the MAGs contained more than 100 RPKM of the genes related to hydrocarbon degradation. Liu et al. [35] also observed multiple copies of HDGs within a single bacterial strain, suggesting that each copy may serve a unique function (e.g., diverse activation patterns). Moreover, we found a statistically significant, yet moderately positive, linear relationship between genome size and HDG count for all MAGs. Benthic MAGs had higher HDG annotations compared to pelagic MAGs and showed a stronger relationship with HDG count compared to the pelagic samples.

The species with the most annotation, *UBA1847 sp022866645* (Pseudomonadota, Woeseiaceae), was found exclusively in benthic samples, followed by *D2472 sp002358345* (Pseudomonadota, D2472), and *MAG-120802 sp030730425* (Actinomycetota, Nanopelagicaceae). On the other hand, *UBA9145 sp001438145* (Pseudomonadota, Pseudohongiellaceae) was only found in pelagic samples. Previous reports identified *UBA1847 sp.* as one of the most abundant bacteria in coastal sediments of the Baltic Sea [69]. This species exhibits a wide range of functions, from facultative chemolithoautotrophy to obligate chemoorganoheterotrophy [70]. The MAGs identified as *UBA1847 sp.* had an average HDG annotation of 39 ± 7 RPKM, suggesting that the most prevalent bacteria in the

benthic ecosystem of the Baltic Sea have the potential to degrade various hydrocarbon substrates. We also recovered MAGs identified as *JAJDAM01 sp.* (Myxococcota A, SMWR01) and *CALGIA01 sp.* (Pseudomonadota, MPNO01) that had the topmost HDG annotations (> 300 RPKM).

Additionally, 66 of the 80 archaeal MAGs we recovered had HDG annotations of more than 10 RPKM. These hydrocarbon-degrading archaea, mostly *Nitrosopumilus sp.* (Thermoproteota, Nitrosopumilaceae), are found in both benthic and pelagic environments. These archaeal species are considered one of the most abundant and ubiquitous microorganisms, primarily found in open oceans and marine sediments [61]. The previous global metagenomic profiling of hydrocarbon-degrading microorganisms found no archaeal genomes [35], whereas assessments in the high Arctic reported three dominant archaeal phyla, i.e., Halobacterota, Euryarchaeota, and Thermoplasmatota. We also identified these phyla in our pelagic samples with low representation. Archaea have fewer genes for hydrocarbon degradation than bacteria [54], so the limited identification of archaeal hydrocarbon degraders might stem from the small pool of sequenced archaeal genomes or the significant phylogenetic divergence between archaeal hydrocarbon degradation genes and the mainly bacterial sequences that have been experimentally validated. The identified archaea and bacteria may be key contributors to hydrocarbon degradation in the benthic and pelagic ecosystems of the Baltic Sea, playing a crucial role in the natural bioremediation process in the area. Moreover, their prevalence and activity suggest an intrinsic mechanism within the benthic and pelagic ecosystems to mitigate pollution, which could have implications for environmental management strategies in the Baltic Sea and similar marine ecosystems.

### HDG distribution showed distinct metabolic adaptations between pelagic and benthic communities

Among the degradation pathways, aerobic metabolism was the most prominent in the Baltic Sea, with HDGs associated with aerobic alkane metabolism being significantly more abundant in pelagic MAGs, reflecting the oxygen-rich conditions of pelagic environments, which favor aerobic processes [71]. On the other hand, HDGs linked to anaerobic alkane metabolism were more prevalent in benthic MAGs, aligning with the low-oxygen or anoxic conditions typically found in seabed sediments [72]. Similarly, Góngora et al. [34] evaluated the hydrocarbon degradation potential of Canadian high Arctic beaches and discovered that alkane metabolism is the prevalent form of hydrocarbon degradation in tidal beach ecosystems. For the aromatic degradation pathways, HDGs involved in anaerobic metabolism were higher in benthic compared to pelagic, while HDGs for aerobic metabolism exhibited similar levels. These findings highlight the ecological specialization of microbial communities in various marine habitats, which is driven by the availability of oxygen and other environmental factors.

Petroleum hydrocarbons are recalcitrant compounds that persist in the environment, influencing microbial communities through prolonged exposure [73,74]. Hydrocarbons in dispersed oil can degrade in aerobic marine waters within days to months, whereas oil on shorelines is often more concentrated, nutrient-limited, and persists in the environment for a longer period [75]. Benthic habitats, characterized by more stable sediments, may experience slower degradation due to limited oxygen and nutrients.^76,77^ In contrast, pelagic environments, characterized by dynamic conditions and enhanced oxygenation, facilitate faster degradation [78]. Consequently, microbial degradation capacities are expected to vary between benthic and pelagic habitats due to differences in microbial communities and physicochemical conditions [79]. The pelagic ecosystem, characterized by its open water column, supports greater microbial and genetic diversity than the benthic ecosystem, which is influenced by sedimentary conditions [80]. The observed higher species and HDG diversity in the Baltic Sea’s pelagic samples may be attributed to environmental factors, e.g., greater oxygen availability, dynamic nutrient cycling, and the presence of diverse organic matter sources [81]. In contrast, the benthic ecosystem often has limited oxygen levels and a more stable, yet nutrient-constrained, environment. These conditions necessitate specific metabolic adaptations among microbial communities, e.g., the ability to utilize alternative electron acceptors or degrade complex organic compounds [82].

Miettinen et al. [33] also explored the hydrocarbon degradation capabilities of bacterial, archaeal, and fungal microbiomes in the Baltic Sea by comparing coastal sites with varying levels of long-term oil exposure. Through the identification of specific genes (i.e., *alkB* and PAH ring-hydroxylating dioxygenase), their study revealed differences in microbial composition and degradation potential between water and sediment samples, as determined by taxonomic profiling using amplicon sequencing. Similar to our observations, the differences in hydrocarbon degradation pathways observed between these ecosystems further underscore the metabolic specialization of microorganisms in different habitats, with certain pathways being more prominent in oxygen-rich pelagic zones and others adapted to the anoxic and suboxic conditions of the benthic ecosystem across the Baltic Sea. This highlights the influence of distinct ecological niches and the possible metabolic adaptations of microbial inhabitants in the benthic and pelagic environments, alongside the intricate interplay between environmental gradients, microbial diversity, and metabolic functions that shape the ecological roles of microorganisms. Moreover, the observed differences between benthic and pelagic microorganisms contribute to their survival and play a crucial role in biogeochemical processes, including nutrient cycling and the degradation of pollutants.

### Long-chain alkanes and dibenzothiophene are the preferred substrates of hydrocarbon degraders

As shown in **Fig. 4d**, the genes associated with the degradation of long-chain alkanes (i.e., C20-C32) and dibenzothiophene were the most prevalent and exhibited the highest abundance. Long-chain alkanes are challenging to degrade due to their large molecular size and hydrophobic nature. Both benthic and pelagic ecosystems contained alkane monooxygenases (*alma Group I* and *III*), which are reported to play crucial roles in the metabolic pathway of extremely hydrophobic long-chain n-alkanes (>C20) that are particularly difficult to degrade [83]. These flavin-binding monooxygenases utilize flavin cofactors to catalyze the oxidation of long-chain hydrocarbons. Additionally, we identified the long-chain alkane monooxygenase (*ladB*) in the Baltic Sea samples, which specializes in oxidizing long-chain alkanes. The dibenzothiophene monooxygenase (dszC) gene was the second most abundant hydrocarbon degradation-associated gene in the Baltic Sea samples. This gene plays a key role in the microbial desulfurization of dibenzothiophenes (DBT), a sulfur-containing compound commonly found in crude oil [84]. As one of the four enzymes in the 4S pathway, *dszC* catalyzes the initial step of converting DBT into DBT sulfone, setting the initial stage for subsequent enzymatic reactions that ultimately remove sulfur without breaking carbon-carbon bonds [85]. Its abundance suggests that the microbial communities of the Baltic Sea are well-suited to handle sulfur-rich hydrocarbons, potentially due to the levels of oil resulting from anthropogenic activities or natural seepage in the region [86]. The presence of these genes reflects the adaptive capabilities of Baltic Sea microorganisms in addressing recalcitrant hydrocarbons, converting them into more reactive and degradable intermediates. This step is crucial for the complete elimination of these hydrocarbons from the environment. Our findings suggest that both benthic sediments and the pelagic water column are equipped to handle hydrocarbon pollution, including oil spills, by leveraging these enzymatic pathways.

### Depth and oil spill volume were the most influential factors on HDG composition

The chemical complexity of oil and its pollution, combined with environmental factors, i.e., depth, oxygen levels, temperature, nutrient availability, and other physical and chemical influences, governs the assembly of microbial communities and the degradation processes they carry out in seawater and marine sediments following oil spills [87]. In particular, we report that depth plays a key role in shaping the composition of genes associated with hydrocarbon degradation in the Baltic Sea. Vigneron et al. [88] reported similar observations, profiling the distribution of microorganisms with hydrocarbon degradation potential at various depths in Lake A in the Canadian High Arctic. In deeper waters, reduced oxygen levels may favor anaerobic degradation pathways, while aerobic processes might dominate in shallower areas [80]. Additionally, sedimentation rates and the accumulation of hydrocarbons in benthic zones can vary with depth, influencing the availability of substrates for microbial degradation [82,88].

Natural hydrocarbon degraders in the Baltic Sea are closely linked to the annual average oil spill documented in each subbasin. For example, the Gulf of Finland, which had the highest annual average oil spill volume record, showed relatively lower HDG diversity (**Supplementary Fig. S5**), having genes associated with the degradation of long-chain (C20-C32) hydrocarbons more prevalent. This suggests the potential enrichment of specific microbial hydrocarbon degraders in the presence of their preferred substrates. On the other hand, the Gulf of Riga and the Quark, where no significant oil spills were recorded, had more diverse HDG profiles. Similar observations were reported for petroleum hydrocarbon-polluted soil, where unpolluted soils had higher diversity and evenly distributed communities than contaminated samples [89]. However, this is not consistent with other subbasins. For example, the Bothnian Bay experienced a low annual oil spill volume and exhibited the highest benthic HDG diversity. Still, these observations indicate that the natural presence of hydrocarbon degraders in the Baltic Sea highlights the importance of understanding spatial variability in microbial responses to oil spills and the corresponding spill volumes. Moreover, salinity and latitude also play significant roles, suggesting that a combination of abiotic factors influences the distribution and activity of these microbial communities. Overall, these observations underscore the significance of depth and salinity as key drivers of microbial responses to oil pollution. They also highlight the spatial heterogeneity of oil spills and their ecological implications.

### Implications for the bioremediation of petroleum hydrocarbons

The volume of oil spills in the Baltic Sea is influenced by a combination of anthropogenic activities, environmental conditions, and governance challenges [13,15,90]. Key factors include the high volume of oil transportation through the region, with major shipping routes and oil terminals contributing to accidental and operational spills [15]. For instance, the Gulf of Finland experiences higher spill levels due to its role as a critical transit area for oil exports [91]. Similarly, the Northern Baltic Proper and Kattegat are busy maritime zones, where dense traffic increases the likelihood of spills [92]. The natural presence of hydrocarbon-degrading microorganisms in the Baltic Sea, coupled with their capacity to adapt to varying oil spill thresholds, can be strategically utilized to estimate the persistence of petroleum hydrocarbons and inform oil spill management. Understanding the functional potential of microbial communities can be used to predict how long petroleum-based pollutants may persist in the environment and how they might be transported across different areas [28,29]. This predictive ability is crucial for guiding cleanup operations, allocating resources more effectively, and prioritizing areas in most urgent need of intervention. The ability to monitor variations in hydrocarbon degradation across regions also supports more precise environmental monitoring. These insights could further inform policies and guidelines for managing oil spills, integrating microbial degradation as a key element of broader mitigation strategies.

Biostimulation techniques can enhance microbial degradation in subbasins with high spill volumes by optimizing environmental conditions like nutrient and oxygen levels [96]. Previous studies have shown that nutrient amendments to contaminated areas, e.g., the Labrador Sea [94], or from potentially at-risk areas, e.g., high-Arctic beaches [95], enhanced the hydrocarbon degradation activity of their native microbial communities. Bioaugmentation, the introduction of robust hydrocarbon-degrading strains in contaminated areas, can also be utilized for cleanup efforts [96]. In less-exposed regions, the resilience of native microorganisms ensures that they can still respond effectively to occasional spills.

## Conclusions

Our study highlights the significant differences in degradation potential between benthic and pelagic environments, as well as among various subbasins within the Baltic Sea. Understanding how environmental factors, i.e., depth and salinity, influence microbial activity enables the development of tailored, site-specific bioremediation plans. By harnessing these microbial communities, oil spill management can become more efficient and sustainable, minimizing environmental damage while supporting ecosystem recovery. Our findings not only enhance our understanding of hydrocarbon-degrading microbial communities in the benthic and pelagic ecosystems of the Baltic Sea but also offer insights into potential microbial-based strategies for the environmental remediation of petroleum hydrocarbons in marine environments. With this, we recommend further assessment and validation of the functional capacities of native microbial communities in the Baltic Sea ecosystem using culture-dependent methods and multi-omic approaches to provide a comprehensive understanding of the specific mechanisms of natural hydrocarbon attenuation in marine ecosystems.

## List of abbreviations

dbRDA: distance-based redundancy analysis;
HDG: hydrocarbon degradation gene;
HELCOM: The Baltic Marine Environment Protection Commission;
HMM: hidden Markov model;
HOLAS: HELCOM holistic assessments;
NMDS: non-metric multidimensional scale.

## Supplementary Information

**Additional file 1** (Supplementary-Information.doc): **Supplementary Fig. S1** Map of the sampling locations of the 203 Baltic Sea metagenomes. Colors indicate the distribution of each sample based on the Baltic Sea sub-basin category of HELCOM (2022). **Supplementary Fig. S2** Histograms showing the distribution of MAG characteristics. (a) Genome size, (b) GC Content, (c) N50 (contigs), and (d) predicted gene counts. Colors represent the environment where the MAGs were located, i.e., pelagic (purple) and benthic (brown). The dashed line indicates mean values. **Supplementary Fig. S3** Scatter plots of the relationship between HDG Count and Genome Size (bp) for (a) all MAGs, (b) benthic MAGs, and (c) pelagic MAGs. **Supplementary Fig. S4** Frequency of HDGs annotated per Baltic Sea sub-basin. **Supplementary Fig. S5** Alpha diversity estimates of the hydrocarbon degradation genes (HDGs) per sub-basin for (a) benthic and (b) pelagic samples. **Supplementary Fig. S6** Upset plot of the shared and unique HMM annotations per Baltic Sea sub-basin. **Supplementary Fig. S7** Maximum-likelihood phylogenomic tree of the archaeal MAGs. **Supplementary Fig. S8** Relative abundance of the top 10 most abundant species visualized (a) per environment and (b) per Baltic Sea sub-basin. **Supplementary Fig. S9** Upset plot of the shared and unique species per Baltic Sea sub-basin. **Supplementary Fig. S10** Bubble plot of the classified microbial taxa at the phylum-level and the occurrence of hydrocarbon degradation genes (HDGs) in each taxa. **Supplementary Fig. S11** Random-forest classification of the relative abundance HDGs across the environments: (left) HDGs presented in descending order of importance, and the (right) relative abundance of HDGs enriched in each environment. **Supplementary Fig. S12** Random-forest classification of the relative abundance of HDG substrates across the environments: (left) Substrates presented in descending order of importance, and the (right) relative abundance of HDG substrates enriched in each environment.

**Additional file 2** (Supplementary-Tables.xls): **Supplementary Table S1** Metagenomics sample information and environmental factors. Metagenomics sequencing data were obtained from the European Nucleotide Archive (ENA) database: accession numbers PRJEB41834 (Broman et al., 2022; Rodríguez-Gijón et al., 2023), PRJEB22997 (Alneberg et al., 2018), and PRJEB34883 (Alneberg et al., 2020). The environmental parameters, i.e., depth (m), salinity (PSU), and temperature (°C), compiled by Rodríguez-Gijón et al. (2023) were used in this study. **Supplementary Table S2** Metagenomics data processing information: Sequence codes, read processing counts (from quality-filtering to contig assembly), prodigal annotations and binning stat values. The Calgary approach to ANnoTating HYDrocarbon degradation genes (CANT-HYD; Khot et al., 2022) counts highlighted in blue. **Supplementary Table S3** Metagenome-assembled genome (MAG) stats and taxonomic classification based on using GTDB-Tk v2.4.0 with R220 (Chaumeil et al., 2022). **Supplementary Table S4** The Calgary approach to ANnoTating HYDrocarbon degrading enzymes database (CANT-HYD) annotations (Khot et al. 2022). **Supplementary Table S5** Count and annotation table of the annotated hydrocarbon degradation genes (HDGs). Reads per kilobase million (RPKM) absolute counts. **Supplementary Table S6.** The annual average of total oil spills during the assessment period 2016-2021 (m³) values (Supplementary Table S6) from HELCOM (2023b). **Supplementary Table S7** Average metagenome-assembled genome (MAG) quality stats per environment. **Supplementary Table S8** Total metagenome-assembled genome (MAG) counts per environment and relative abundance based on species-level classification. **Supplementary Table S9** Statistical results of the permutation test for homogeneity of multivariate dispersions and permutation test for adonis under a reduced model of the hydrocarbon degradation genes (HDG) dataset. **Supplementary Table S10** Differential abundance test by Random Forests for the HDGs per environment and per subbasin. **Supplementary Table S11** Statistical results of the permutation test for distance-based redundancy analysis (dbRDA) analyses in R of the ARG composition and environmental factors. **Supplementary Table S12** Spearman rank correlations between environmental variables and the RPKM abundance of the hydrocarbon degradation genes (HDG).

## Declarations

### Ethics approval and consent to participate

Not applicable.

### Consent for publication

Not applicable.

### Availability of data and materials

All the raw sequence data are obtained from the European Nucleotide Archive (ENA) database with project accession numbers PRJEB41834 (Broman et al., 2022; Rodríguez-Gijón et al., 2023), PRJEB22997 (Alneberg et al., 2018), and PRJEB34883 (Alneberg et al., 2020). The specific accession numbers for each metagenome are listed in Table S1. Additional data from the analyses presented in this paper are available in the Supplementary Figures and Tables. The corresponding visualization and downstream analysis codes are publicly accessible at https://github.com/jserrana/baltic-hdg.

### Authors’ contributions

JMS, BD, EB, and MP conceptualized the study. BD compiled the HELCOM literature and data. EB and FN provided support on the bioinformatics framework and analysis. JMS performed data analysis and wrote the first draft of the manuscript. All authors contributed to the writing and editing of the manuscript.

## Acknowledgments

The authors acknowledge the National Academic Infrastructure for Super-computing in Sweden for granting us access and storage to the Uppsala Multidisciplinary Center for Advanced Computational Science (UPPMAX) computational infrastructure.

