## Supplementary Information for "Microbial hydrocarbon degradation potential of the Baltic Sea ecosystem"

*Authors & Affiliations*

Joeselle M. Serrana[1,2]*, Benoît Dessirier[1,3], Francisco J. A. Nascimento[1,4], Elias Broman[1,3,4], and Malte Posselt[1,2]

[1] Stockholm University Center for Circular and Sustainable Systems (SUCCeSS), Stockholm University, 106 91 Stockholm, Sweden
[2] Department of Environmental Science (ACES), Stockholm University, 106 91 Stockholm, Sweden
[3] Baltic Sea Centre, Stockholm University, Stockholm, Sweden
[4] Department of Ecology, Environment, and Plant Sciences (DEEP), Stockholm University, 106 91 Stockholm, Sweden

**Supplementary Table S12.** Spearman rank correlations between environmental variables and the RPKM abundance of the hydrocarbon degradation genes (HDG).

### Supplementary Figures

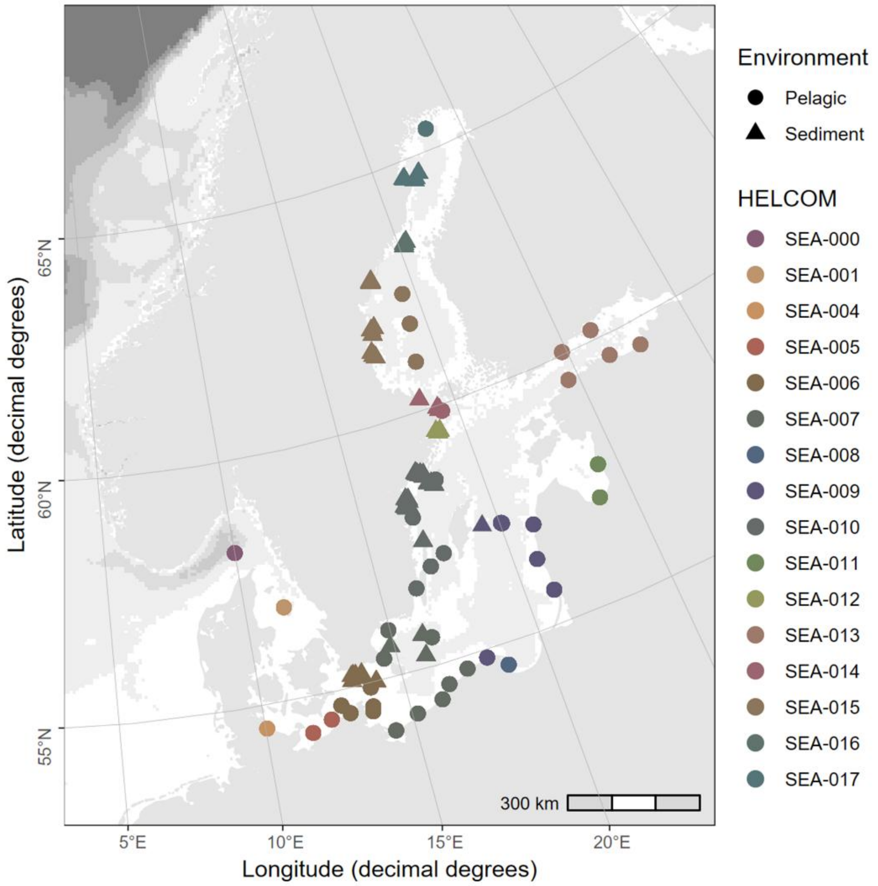


**Supplementary Fig. S1. Map of the sampling locations of the 203 Baltic Sea metagenomes.** Colors indicate the distribution of each sample based on the Baltic Sea sub-basin category of HELCOM (2022).

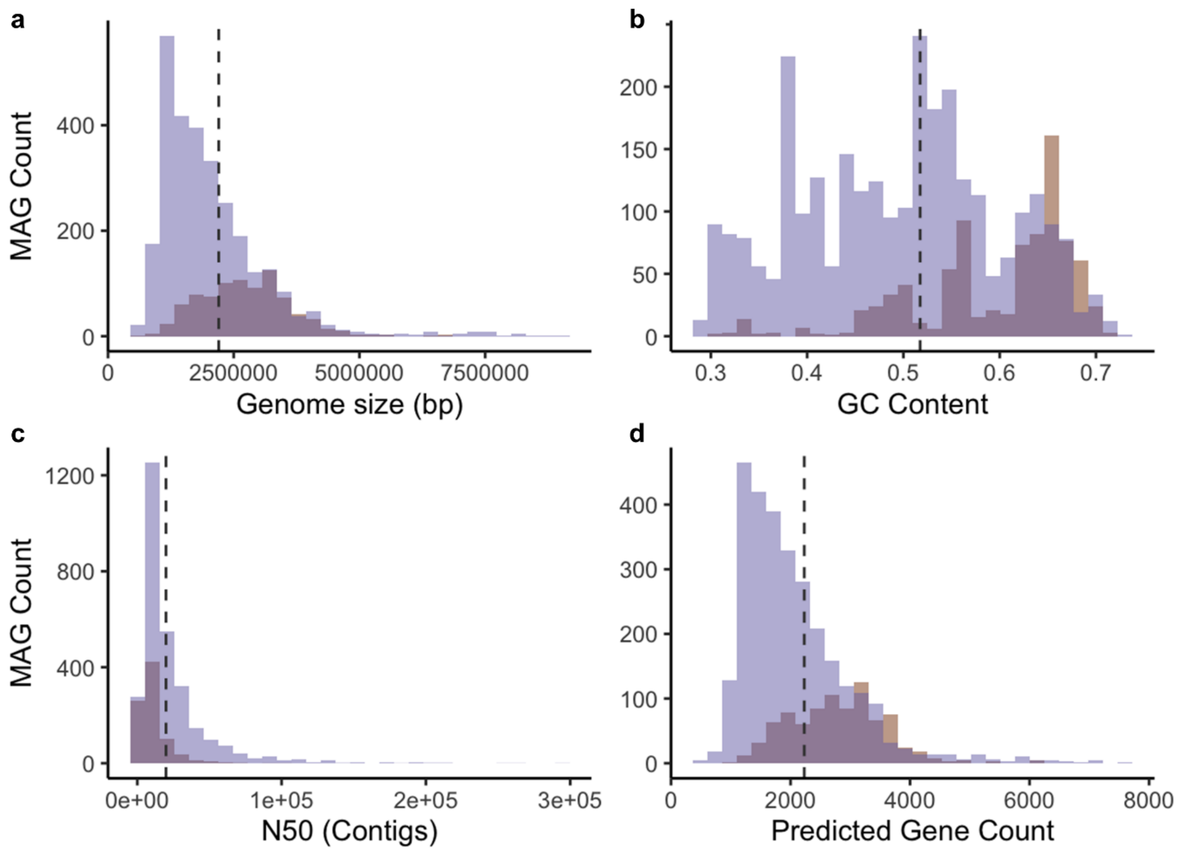


**Supplementary Fig. S2. Histograms showing the distribution of MAG characteristics.** (a) Genome size, (b) GC Content, (c) N50 (contigs), and (d) predicted gene counts. Colors represent the environment where the MAGs were located, i.e., pelagic (purple) and benthic (brown). The dashed line indicates mean values.

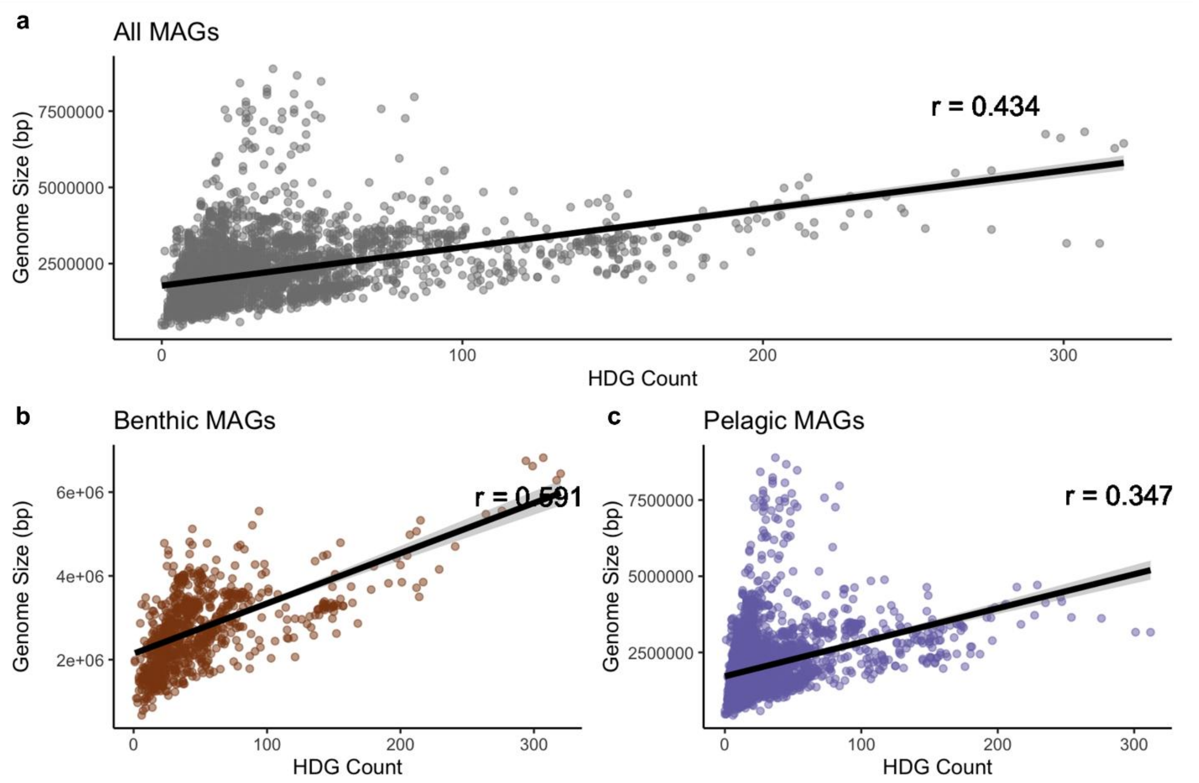


**Supplementary Fig. S3.** Scatter plots of the relationship between HDG Count and Genome Size (bp) for (a) all MAGs, (b) benthic MAGs, and (c) pelagic MAGs.

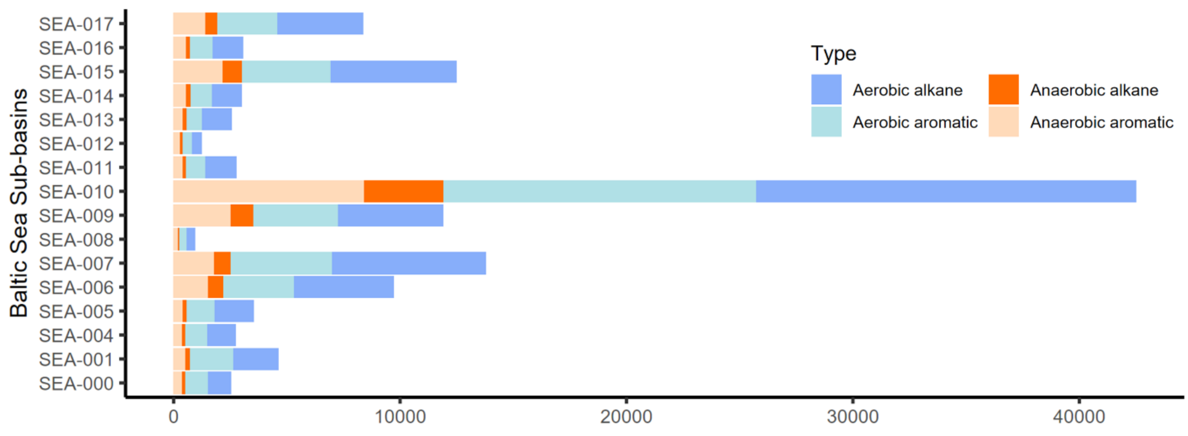


**Supplementary Fig. S4.** Frequency of HDGs annotated per Baltic Sea sub-basin**.**

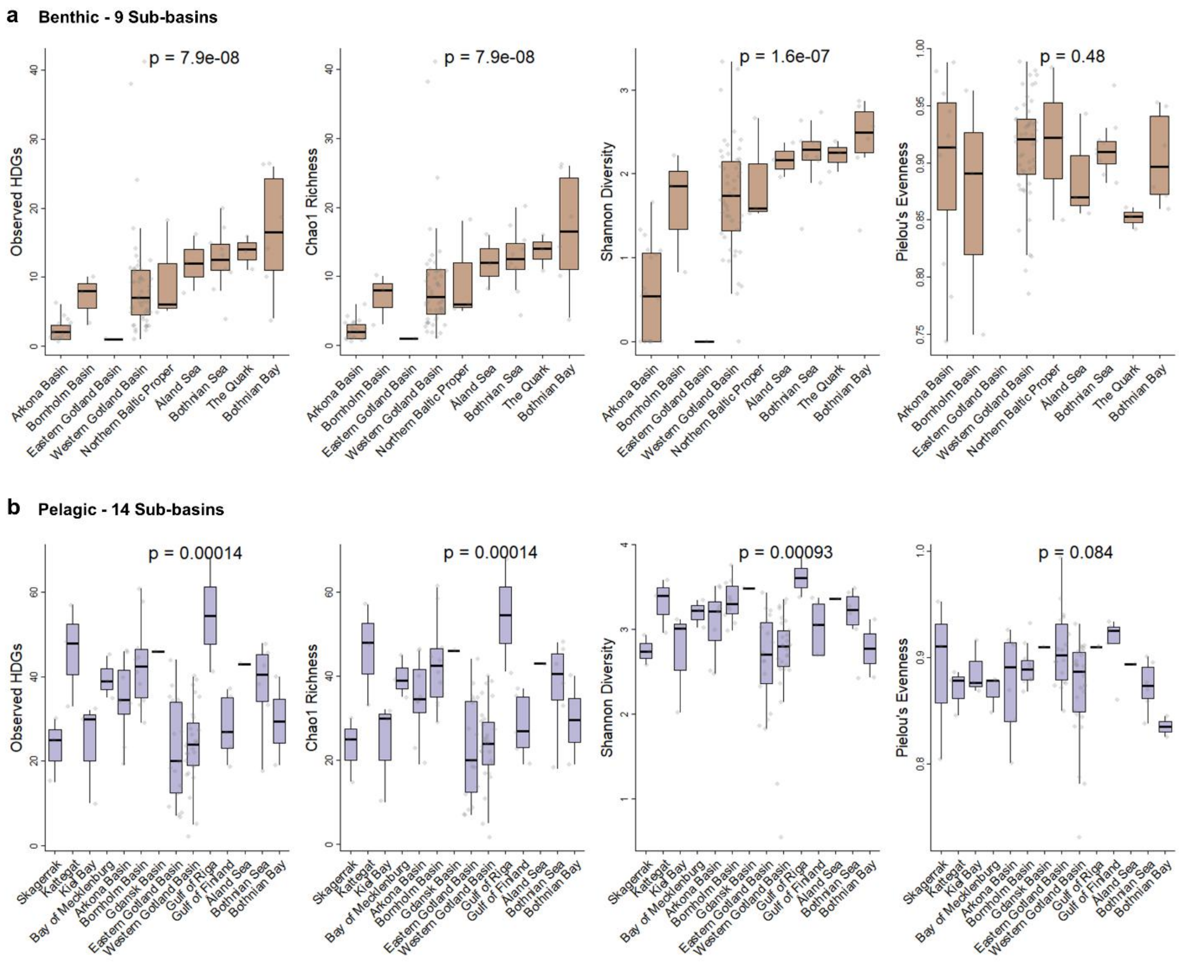


**Supplementary Fig. S5.** Alpha diversity estimates of the hydrocarbon degradation genes (HDGs) per sub-basin for (a) benthic and (b) pelagic samples.

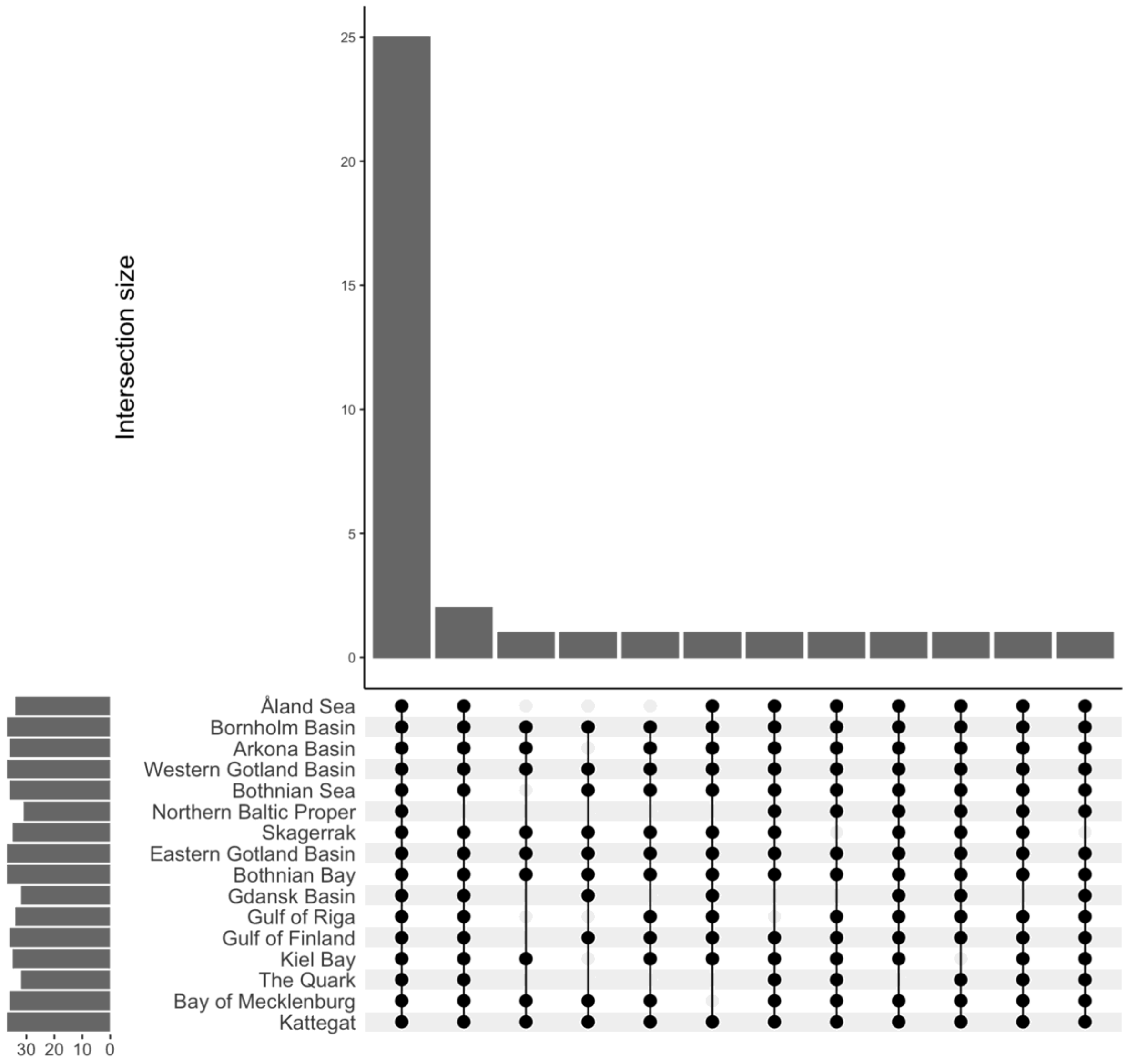


**Supplementary Fig. S6.** Upset plot of the shared and unique HMM annotations per Baltic Sea sub-basin.

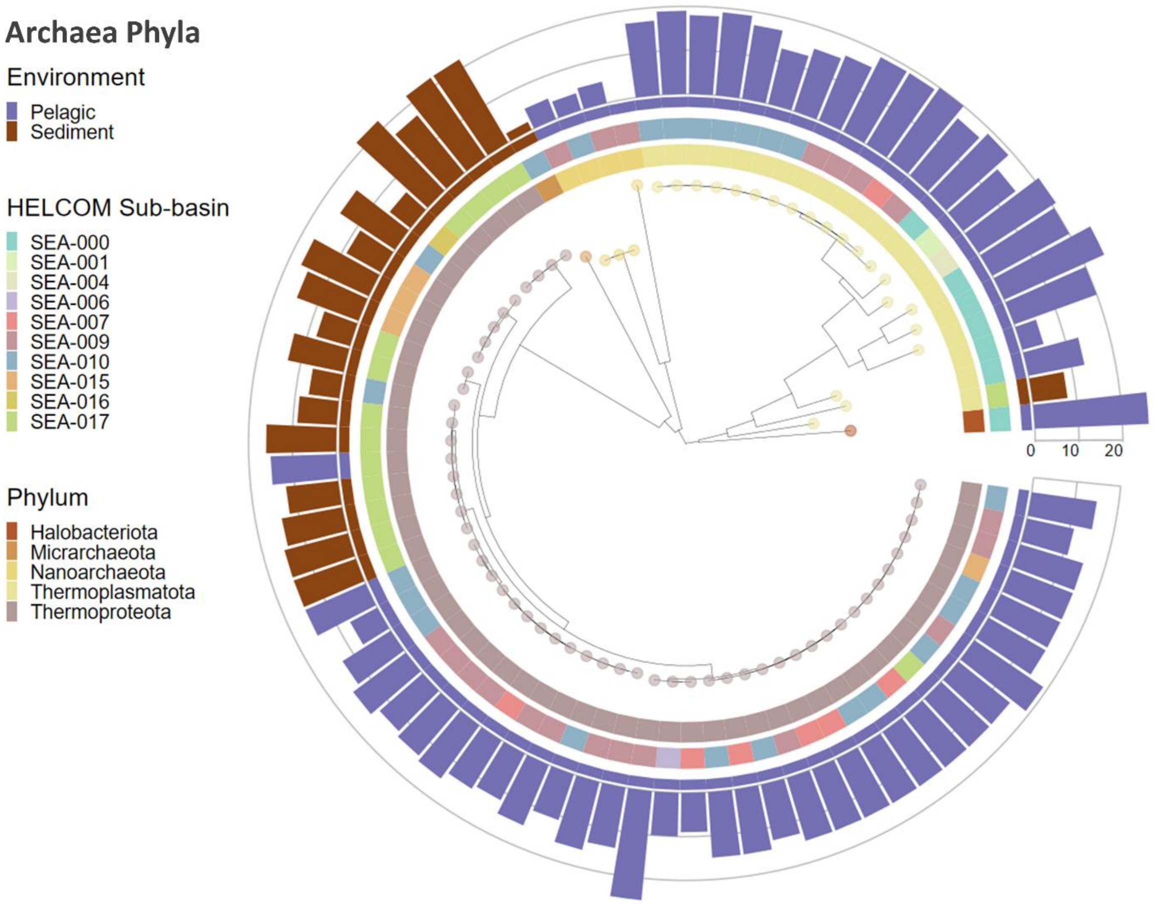


68

69 **Supplementary Fig. S7.** Maximum-likelihood phylogenomic tree of the archaeal MAGs.

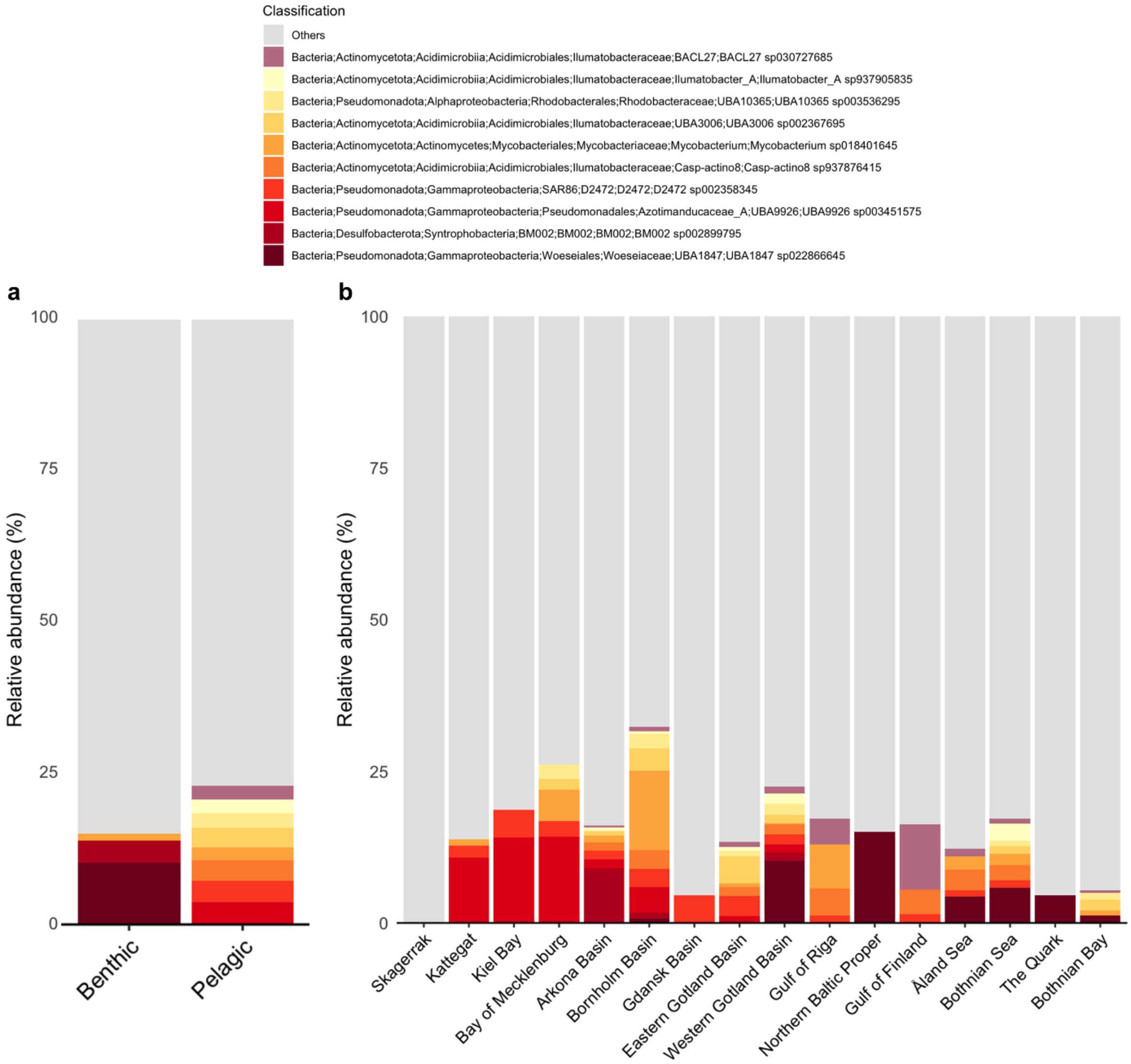


**Supplementary Fig. S8.** Relative abundance of the top 10 most abundant species visualized (a) per environment and (b) per Baltic Sea sub-basin.

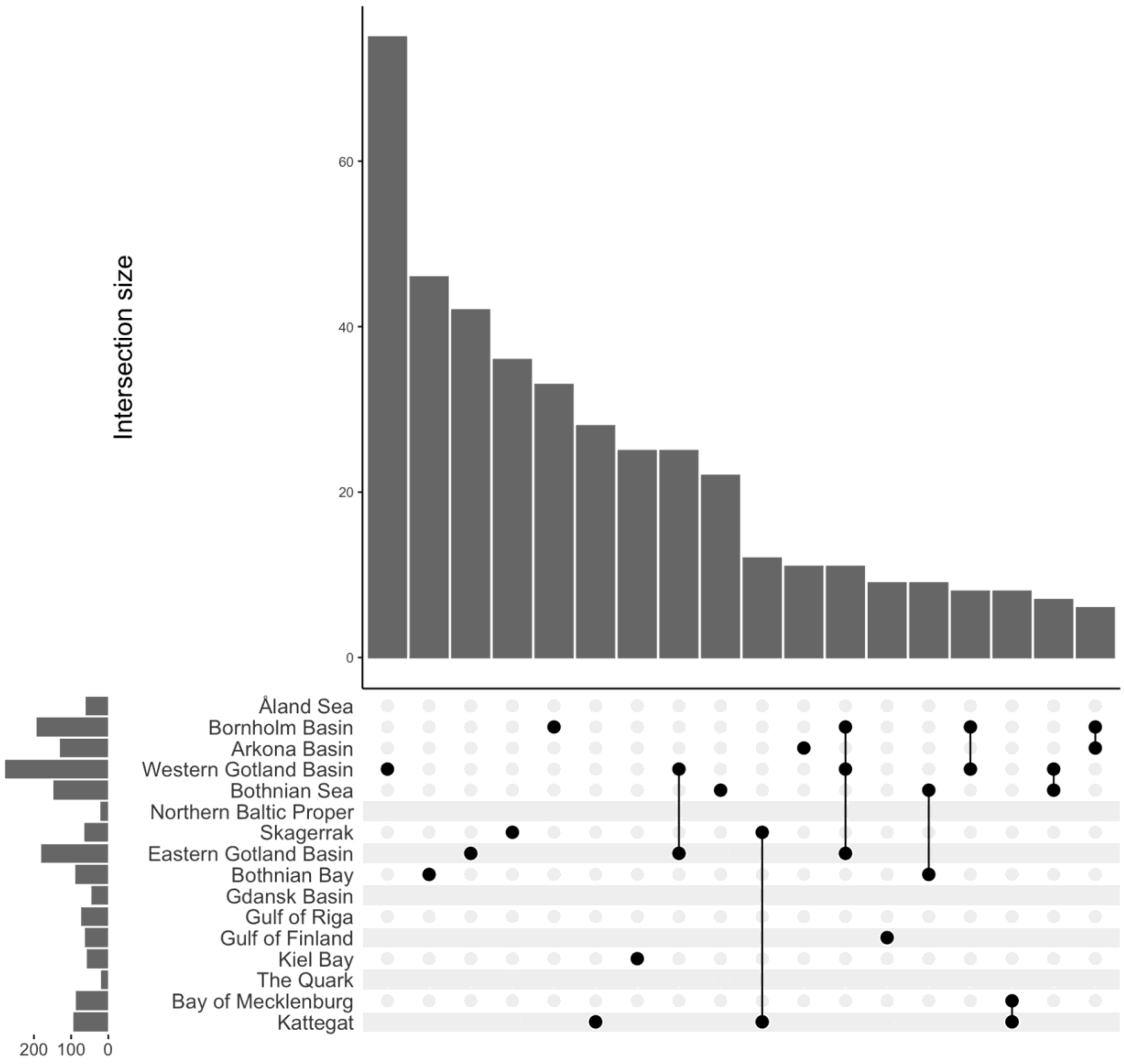


**Supplementary Fig. S9.** Upset plot of the shared and unique species per Baltic Sea sub-basin.

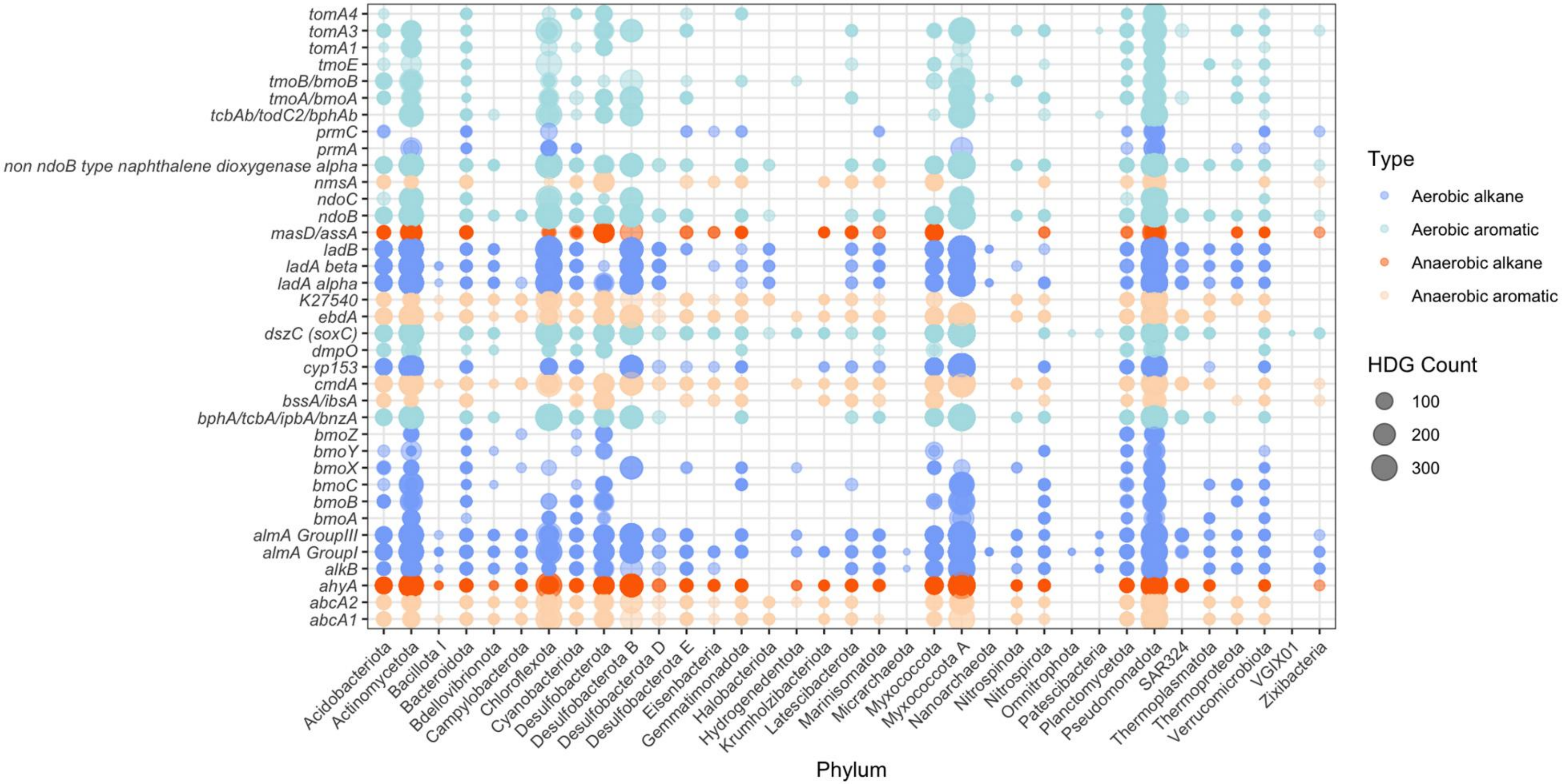


**Supplementary Fig. S10.** Bubble plot of the classified microbial taxa at the phylum-level and the occurrence of hydrocarbon degradation genes (HDGs) in each taxa.

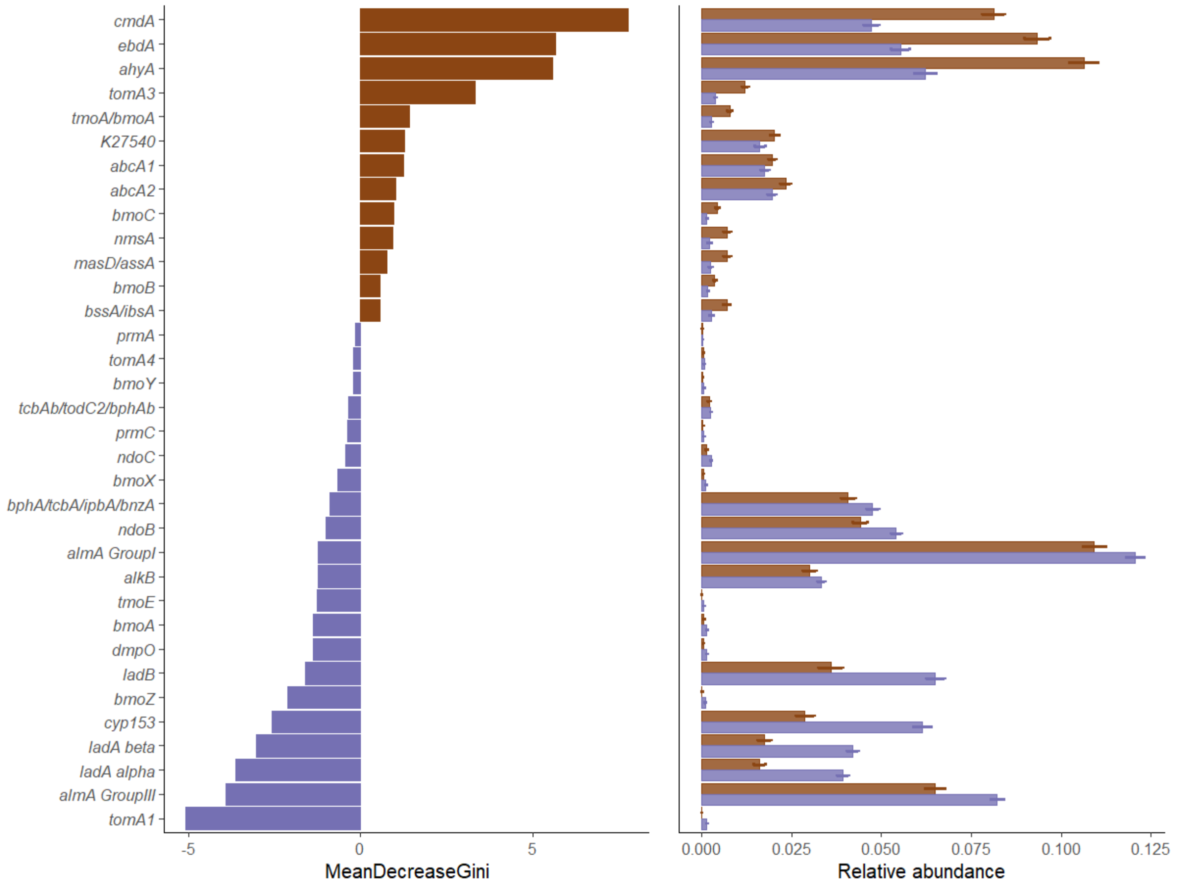


**Supplementary Fig. S11.** Random-forest classification of the relative abundance HDGs across the environments: (left) HDGs presented in descending order of importance, and the (right) relative abundance of HDGs enriched in each environment.

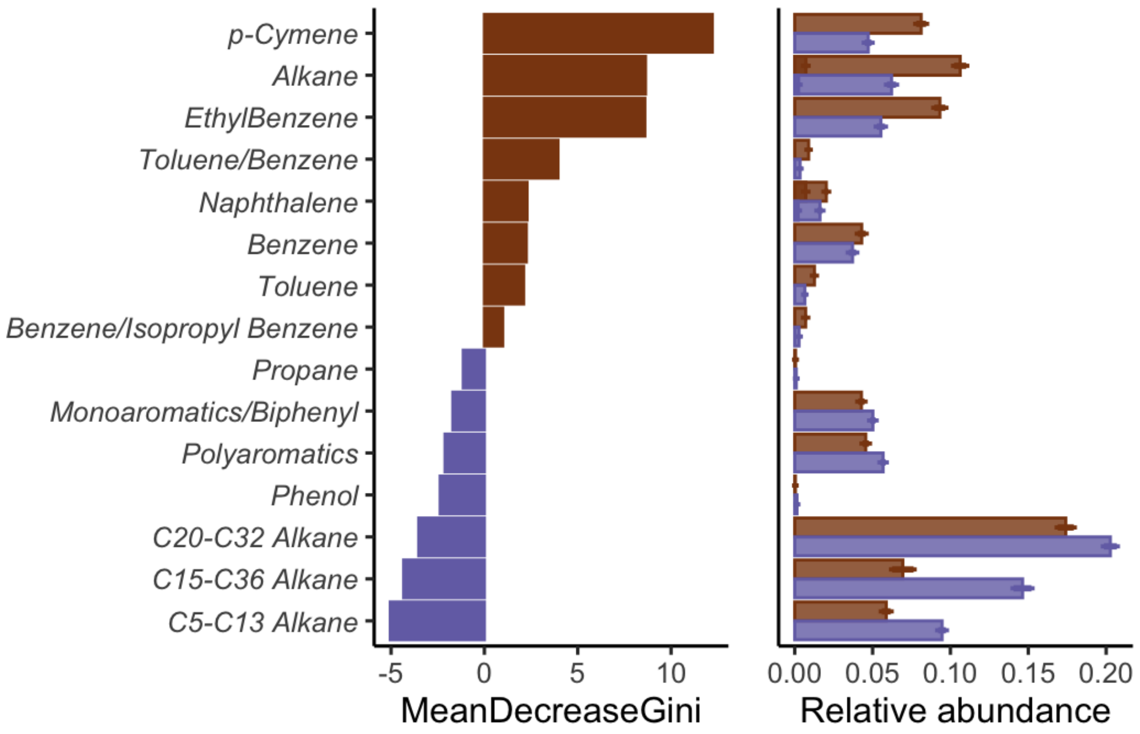


**Supplementary Fig. S12.** Random-forest classification of the relative abundance of HDG substrates across the environments: (left) Substrates presented in descending order of importance, and the (right) relative abundance of HDG substrates enriched in each environment.
